# Cellular coordination underpins rapid reversals in gliding filamentous cyanobacteria and its loss results in plectonemes

**DOI:** 10.1101/2024.02.06.579126

**Authors:** Jerko Rosko, Kelsey Cremin, Emanuele Locatelli, Rebecca N. Poon, Mary Coates, Sarah J. N. Duxbury, Kieran Randall, Katie Croft, Chantal Valeriani, Marco Polin, Orkun S. Soyer

**Author notes:** These authors contributed equally (Jerko Rosko, Kelsey Cremin, Emanuele Locatelli, Rebecca N. Poon). Medical Research Council, Weatherall Institute of Molecular Medicine, University of Oxford (Katie Croft).

## Abstract

2

Cyanobacteria are key contributors to biogeochemical cycles through photosynthesis and carbon fixation. In filamentous, multicellular cyanobacteria these functions can be influenced through gliding motility, which enables filaments to localise in response to light and also form aggregates. Here, we use the aggregate forming species *Fluctiforma draycotensis* to study gliding motility dynamics in detail. We find that filaments move in curved and straight trajectories interspersed with re-orientation or reversal of direction. Most reversals take few seconds but some take substantially longer, resulting in a long-tailed distribution of stoppage times. Mean filament speeds range around a micron per second with a relatively uniform distribution against filament length, implying that all or fixed proportion of cells in a filament contribute to movement. We implement a biophysical model that can recapitulate these findings. Model simulations show that for filaments to reverse quickly, cells in a filament must achieve high coordination of the direction of the forces that they generate. To seek experimental support of this prediction, we track individual cells in a filament. This reveals that cells’ translational movement is fully coupled with their rotation along the long-axis of the filament, and that cellular movement remains coordinated throughout a reversal. For some filaments, especially longer ones, however, we also find that cellular coordination can be lost, and filaments can form buckles that can twist around themselves, resulting in plectonemes. The experimental findings and the biophysical model presented here will inform future studies of individual and collective filament movement.

**Significance Statement:** Cyanobacteria contribute to global oxygen production and carbon capture. Some cyanobacteria exist as multicellular filaments and display gliding motility that allows them to respond to light and to form aggregates, which influences their biological functions. Here, we study the dynamics of gliding motility. We find that filaments’ movement is interspersed with re-orientation or reversal of direction and that mean filament speed is mostly independent of filament length. We implement a biophysical model that predicts these features to relate to cells in a filament having high coordination of the direction of the forces that they generate. We find experimental support for this predicted cellular coordination, but also discover instances of longer filaments loosing coordination, resulting in buckling and entangling with other filaments.

## 3 Introduction

Cyanobacteria are key contributors to global primary production and oxygen generation [1, 2]. They display high diversity and are adapted to a wide range of habitats from the soil crust to freshwater, and the ocean [2, 3]. Within this diversity, some cyanobacterial species display multicellularity, existing as filaments composed of multiple individual cells. Many of these filamenteous cyanobacteria display a specific type of motility, known as gliding motility [3, 4, 5, 6]. Gliding motility enables filaments to position themselves in response to light [7, 8, 9, 10] and also to form aggregates [11, 12, 13], thereby enhancing photosynthesis and additional functions such as nitrogen fixation and iron acquisition [14].

Gliding motility is a specific form of bacterial motility that occurs on surfaces without significant deformation or use of flagella [15, 16, 17, 18, 6, 19]. In single celled bacteria, studies in the species *Flavobacterium johnsoniae* and *Myxococcus xanthus* have shown that gliding motility involves membrane-bound protein complexes moving in helical patterns across and around the cell body [20, 21, 22, 23]. These membrane proteins are suggested to reach so-called focal adhesion points, at cell-surface contact points, where they convert their motion into axial forces that translate the cell and rotate it around its long axis [24, 18]. Mutational studies in these species, as well as filamenteous cyanobacteria, have also identified proteins involved in the biosynthesis and secretion of a sugar polymer, so-called slime, to be essential for gliding motility [15, 16, 17, 18, 25, 26]. In filamentous cyanobacteria, helically organised surface fibril proteins [27, 28, 29, 30] and junctional protein complexes [31] were identified and suggested to link with motility and slime secretion [31, 27]. Type IV pili machinery is also implicated in filamenteous cyanobacteria gliding, both in *Nostoc punctiforme*, where only differentiated filaments called hormogonia display gliding motility[4], and in other species [32, 33, 34, 8].

Despite these ongoing efforts on identifying the molecular mechanisms of force generation, the dynamics of gliding motility in filamentous cyanobacteria remains poorly characterised. In particular, it is not clear how filamentous cyanobacteria achieve coordination of cellular propulsive forces during gliding [6]. A better understanding of motility dynamics can provide insights into cellular coordination and inform molecular studies of propulsion. Study of the motility dynamics at the filament level is also relevant to population-level observations, such as aggregate formation. Current theoretical explanations for these observations make use of specific assumptions about filament behaviour [35, 36], which can be better justified, or ruled out, upon better understanding of movement dynamics.

Here, we study gliding motility dynamics in a filamentous cyanobacterium *Fluctiforma draycotensis*, capable of gliding motility and extensive aggregate formation [13]. We focus specifically on characterisation of movement dynamics, rather than elucidating their underlying molecular mechanisms. To this end, we used extensive time lapse microscopy under phase, fluorescent, and total internal reflection (TIRF) modalities. We found that gliding motility involves movement on curved or straight trajectories interspersed with rapid re-orientation or reversal events and with mean speeds independent of filament length. We were able to recapitulate these findings with a physical model of filament movement, which predicts a high importance for cellular coordination to achieve rapid reversals. We were able to confirm this prediction by tracking individual cells during a reversal. The single-cell analysis also revealed a direct coupling of translational and rotational movement, which can generate torsional forces during reversals. In support of such forces, we found that longer filaments tend to readily buckle and twist upon themselves to form plectonemes. The formation of plectonemes is associated with loss of cell-to-cell coordination during reversals, leading to filament ends moving independently. The presented findings quantify individual filament movement and provide a physical model of gliding motility that is consistent with experimental observations. Together, they will inform future studies on the molecular and physical mechanisms of force generation and collective behaviour of filaments.

## 4 Results

### Filament movement is interspersed with stoppage events, whose duration has a long-tailed distribution

Using time-lapse microscopy, we observed filament movement on glass slides and under agar pads (see *Methods*). We found that filaments move on straight or curved trajectories that are interspersed with re-orientation or reversal of direction (see Supplementary Movies S 1 and S 2). Under agar, it was possible to distinguish a region of different phase contrast, which aligned with the trajectories of the filaments, forming a ‘track’ (Fig. 1A and Movie S 3) (we note that it is possible that the tracks are associated with the secreted slime, see *Discussion* section). Lone filaments’ trajectories were mostly confined to these tracks, involving a stoppage and reversal at each end of the track (Fig. 1B), while filaments entering a circular track did not readily reverse (Movie S 4). We also observed filaments sometimes reversing without reaching track ends, but when this happened, it was usually at a consistent location (Movie S 5). Thus, the observed tracks under phase microscopy constitute a confining space that bounds filament movement and dictates reversal frequency (see Supplementary Fig. S1). On glass, filament movement still displayed a go-stop-go pattern, but stoppages led to both reversal and re-orientation of direction, and as a result, trajectories were less well-defined (Movie S 2). Reversals and re-orientations were associated with a deceleration-acceleration of filaments (Fig. 2A,B and Fig. S2). This decrease in speed can either result from external counter forces or loss of propulsive force, for example through filaments reaching end of a track under agar or detaching from the surface on glass (see Movies S 1 and S 2). Mean filament speeds ranged around one micron per second and displayed a relatively uniform distribution against filament length on agar (Fig. 2C) and a weak positive correlation with filament length on glass (Fig. S3A). Since filament speed results from a balance between propulsive forces and drag, these observations of no or positive correlation between filament speed and length show that all (or a fixed proportion of) cells in a filament contribute to propulsive force generation.

**Figure 1:**
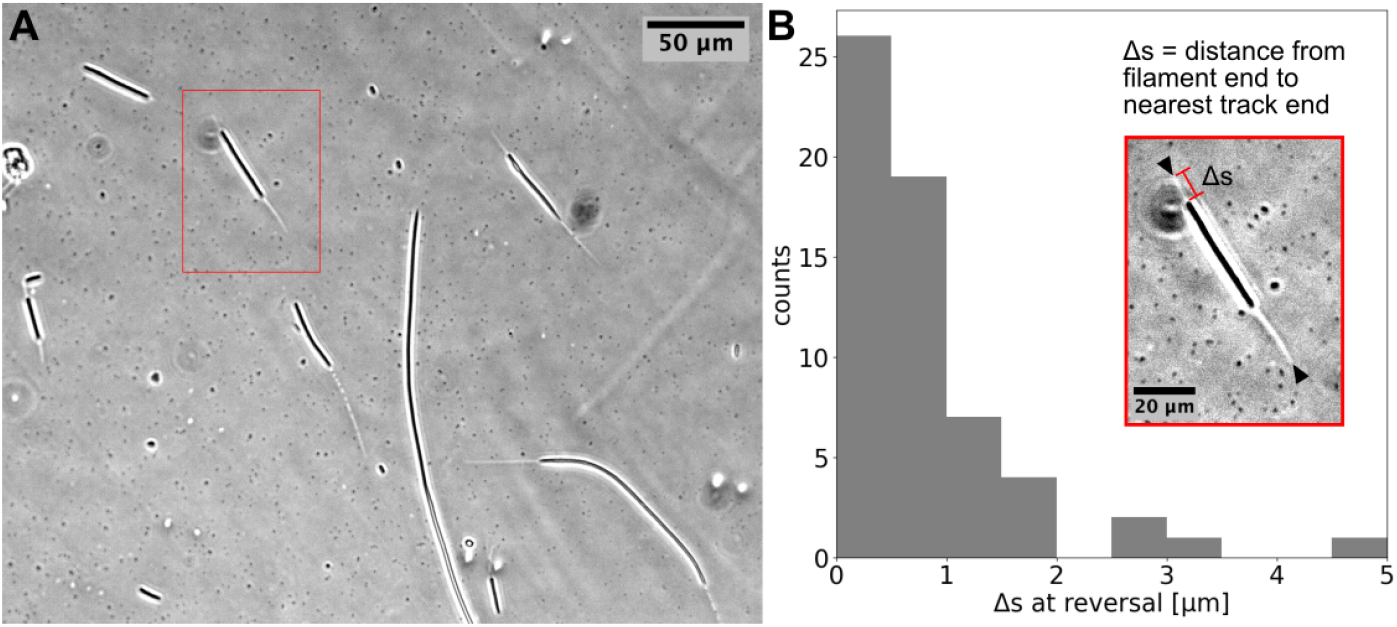
**(A)** Phase contrast image of filaments on an agar pad, showing ‘tracks’ associated with moving filaments. **(B)** Distribution of distance between the end of the filament and the closest track end (Δ*s*) at reversals (see Inset). Data is from multiple filaments (*n* = 14), each showing several reversals, resulting in 61 data points. Approximately 70% of the Δ*s* values are less than the measurement error (≈1 µm). Inset: the single filament and track highlighted in **(A)** with a red box. Track ends are indicated by black arrowheads.

**Figure 2:**
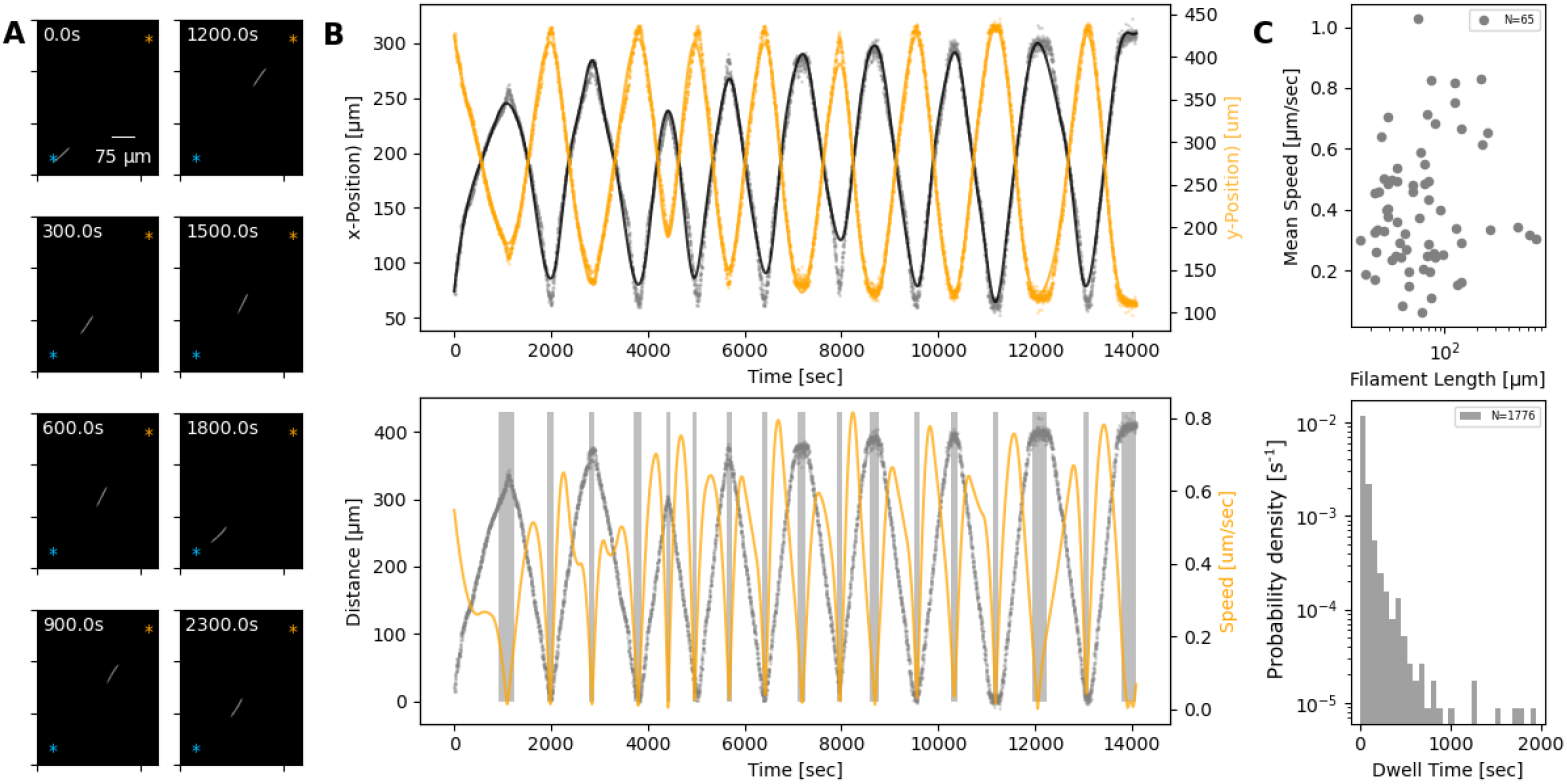
**(A)** A single filament shown at different time points of its movement under an agar pad. Time-lapse images were captured at 2 second intervals using fluorescence microscopy. The scale-bar shown on the first image applies to all subsequent ones. The extreme points of the trajectory across the time lapse are marked with blue and orange asterisks on each image. **(B) Top:** X- and Y-coordinates of filament’s centre throughout the recorded time-lapse. The points show observations, while the line shows a spline fit to this data. **Bottom:** The distance (gray) between filament centre and one of the extreme ends of its trajectory, shown with blue asterisk on panel A, and filament speed (orange) throughout the time-lapse. The speed is calculated from the spline fitted to the x- and y-coordinates shown in the top panel. Gray backdrop regions indicate time-points with speed below a set threshold, indicating reversal events. **(C) Top:** Mean filament speed from 65 different filaments observed under agar, plotted against filament length. **Bottom:** Distribution of dwell times, as calculated from independent reversal events. For the same analyses for observations on glass, see (Fig. S3).

We can consider the reversal / re-orientation events of a filament as akin to tumbling events seen in flagella-based bacterial motility. To this end, we were interested in the distribution of time spent during reversals, which we call the “dwell time”. Analysing over 1700 reversal events across 65 filaments, we found that dwell times display a long-tailed distribution (Fig. 2C). On glass, we found a similar distribution from 1434 reversal / re-orientation events (Fig. S3B). Most reversals happened within few seconds, while a minority involved significant time spent stationary. We found that there was no clear difference between dwell time of leading and trailing ends of the filament (Fig. S5). Taken together, these findings show that filament reversals are mostly rapid events that do not involve delays across the filament. The filaments on agar reverse with a frequency that is inversely proportional to the filament length (which is in turn proportional to the track length) (see Fig. S1). In contrast, we find that the frequencies of reversals on glass do not show a correlation with the filament length and are narrowly distributed (see Fig. S4). These findings are inline with the idea that tracks on agar are defined by filament length, and dictate reversal frequency, resulting in strong correlations between reversal frequency, track length, and filament length. On glass, filament movement is not constrained by tracks, and we have a specific reversal frequency independent of filament length.

### A biophysical model captures filament movement dynamics and highlights a key role for cell-to-cell coordination for achieving fast reversals

As summarised in the introduction, the molecular basis of propulsive forces is not fully understood. However, the presented quantification of motility dynamics highlights three key observations that can be used to develop a plausible biophysical model of filament movement. Firstly, and most obviously, cells in a filament remain physically linked during motion and therefore must exert mechanical forces onto each other. Secondly, all or most cells contribute to propulsive force generation, as seen from a uniform distribution of mean speed across different filament lengths (Fig. 2C and S3). Thirdly, and finally, reversals are associated with reduction in speed (or stoppage) and usually happen within seconds.

We implement the first two observations by modelling cells in a filament as beads connected by springs, with each cell capable of generating propulsion and exerting pull and push forces onto others (see Fig. 3A and *Methods*). The third observation shows that cells have a mechanism to alter the direction of their propulsive force. To this end, we model reversals as resulting from a stochastic cellular signalling pathway that is linked to mechano-sensing of external and internal forces acting on a given cell (see *Methods* and Eqs. (3),(6)). This modelling choice effectively assumes that cells have an intrinsic reversal rate, but adjust this in such a way to reduce any compression or extension that they perceive. In addition, and to account for cellular sensing dynamics, we introduce a memory (i.e. a refractory period) that limits cells ability to switch force direction if they have recently done so (see Eqs. (3),(4) in *Methods*).

**Figure 3:**
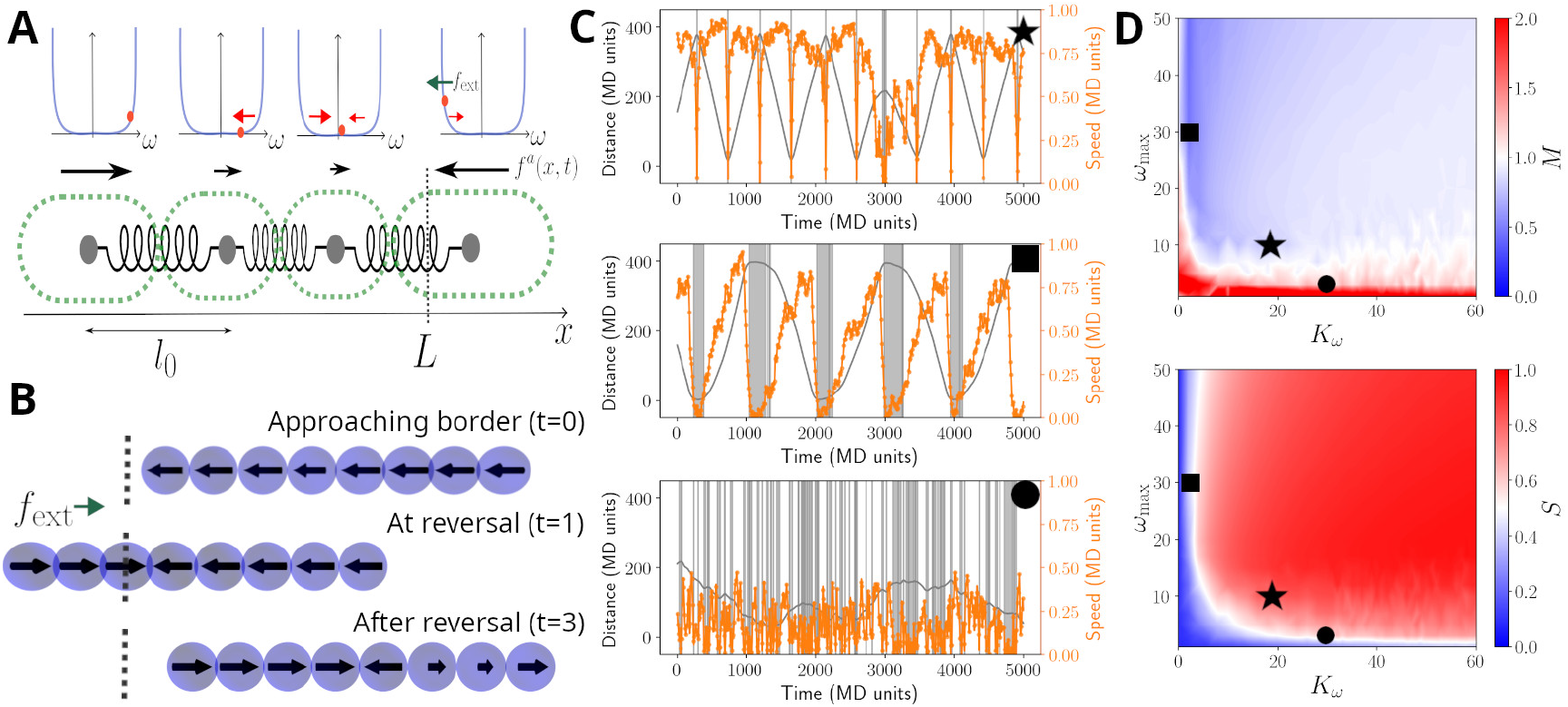
**(A)** A cartoon of the biophysical model (see *Methods*). Cells are modelled as beads, connected by springs, with preferred rest length *l*_0_. Cells self-propel with a propulsion force *f*_*a*_. It is assumed that cells regulate the direction of *f*_*a*_, and this regulation is modelled by a function *ω*, which includes a self-regulatory element, implemented as a confining potential *U*_*ω*_ (shown as blue lines in the panels above cells). The value *ω*_*i*_ of each cell is represented with an orange dot and its affected by random fluctuations, a mechanical feedback from neighbouring cells (*V*_*ω*_, red arrows), and an external signal *f*_ext_, present at the ends of the track *L* (green arrow) (see Eq. (5)). **(B)** Example of reversal process: the snapshots show a coordinated filament approaching the border (top); after reaching it, the closest cells reverse their propulsion under the action of *f*_ext_ (centre). This prompts the rest of the cells to reverse and the filament coordinates again to travel in the opposite direction (bottom). **(C)** Simulated trajectories (gray lines) and absolute speed value (orange lines) as a function of time, presented as in Fig. 2B, for *N*_*f*_ =70 units in a track of length *L/l*_0_ =400. The different panels display a typical trajectory for a well behaved filament (top, *K*_*ω*_ = 20, *ω*_max_ = 30), a filament with little cell-to-cell coupling (centre, *K*_*ω*_ = 1, *ω*_max_ = 30) or with little memory (bottom, *K*_*ω*_ = 30, *ω*_max_ = 1). The gray bands highlight reversal events and their duration. **(D)** Contour plots of the synchronisation *S* (top) and of the reversal efficiency *M* (bottom) as a function of the cell-to-cell coupling *K*_*ω*_ and of the cell memory *ω*_max_. See Methods section for the definition of *S* and *M*. Black symbols highlight the systems showcased in panels **(C)**.

Given this model, we simulated movement of filaments in a 1-dimensional space and in the presence of an external force field. The latter allows us to implement a reduction of speed and analyse filament movement on a defined track, as observed experimentally for the movement under agar (Fig. 2). For appropriate choice of parameters, we found that this model can produce sustained back-and-forth movement (Fig. 3B and Movie S 6). We also run simulations in the absence of an external force field, more closely mimicking the case on glass. Here, again, the model was able to reproduce reversals and experimentally observed dwell time distributions (Fig. S12 and S13).

The ability of the model to present reversals can be understood from the way it implements mechano-sensing at the cell level. On simulations mimicking the agar, when leading cells reach the track ends, the external force field causes an increase in their tendency to reverse the direction of their propulsive force (Fig. 3B). When this happens, trailing cells are compressed by both those in front and behind them, and, if cell-cell coupling is strong enough, are forced to reverse as well. Thus, the mechano-sensing creates a signal that travels across the filament and sustains a coordinated reversal event. The same dynamics can also be created from stochastic reversal of few individual cells, and as such, we also observed reversals in the model simulations without the filament reaching the track ends, or without external force field for some parameter values (Fig. 3B, Fig. S12 and S13).

To better understand how the key model parameters affect reversals, we run simulations across a wide range of parameter values (Fig. 3C,D). To quantify reversal dynamics, we considered two measures that are based on cell-to-cell coordination and extent of reversals. The reversal-related measure, *M*, gives the ratio of the number of actual reversals over the estimated number of reversals obtained by considering an idealised filament, that has a defined mean speed and reverses deterministically and instantaneously at track ends (Eq. (7). Thus, a value of *M* ≈ 1 can indicate regular and rapid reversals at track ends, although it can also indicate simply a high reversal frequency. The coordination measure, *S*, gives the extent of homogeneity of cellular propulsive forces across a filament (Eq. (9)), with a maximum value of 1, corresponding to completely synchronised cell motion. Evaluating *S* and *M* together allows us to better quantify coordination in reversal dynamics along a track. We find that both measures are sensitive to the strength of the cell-to-cell coupling and the memory parameter (Fig. 3C). Indeed, when both parameters are reduced, reversals become more erratic, with filaments undergoing many reversals and individual cells having opposing propulsive force directions (Fig. 3B and Movie S 7). As expected from these results, these two parameters also affect the shape of the dwell time distribution resulting from model simulations (Fig. S6). For sufficiently high cell-to-cell coupling and memory parameters, simulated filaments reverse rapidly, and primarily at track ends, and cellular propulsive force directions remain homogeneous throughout their movement (Fig. 3B). To explore if real filaments’ movement conform to such expected dynamics arising from high cell-to-cell coupling, we quantified the number of expected reversals using individual filament and track lengths, and the observed mean filament speed from all observations. We found that for the high majority of the filaments, the number of observed reversals matches closely with the expected number of reversals, in qualitative agreement with the model simulations performed with sufficiently high cell-to-cell coupling parameter (Fig. S7).

### Filament movement shows high cell-to-cell coordination and coupling of translation with rotation

Taken together, the experimental and model results presented so far show that most reversals are rapid events, and to achieve them the filaments should have high cell-to-cell coupling in their movement. To explore this model-highlighted idea of cellular coordination and to better understand what happens during a reversal at the single cell level, we undertook Total Internal Reflection Fluorescence (TIRF) microscopy, which illuminates only an ≈200 nm thick band of the sample and allows observation of surface features (Fig. 4A-B, top panels). This allowed us to clearly identify individual cells and their septa with neighboring cells, which we tracked to quantify individual cell movement. We found that cell movement across the filament mostly stays coordinated during a reversal, in terms of speed and direction of movement (Fig. 4A, bottom panels, see also associated Movie S 8).

**Figure 4:**
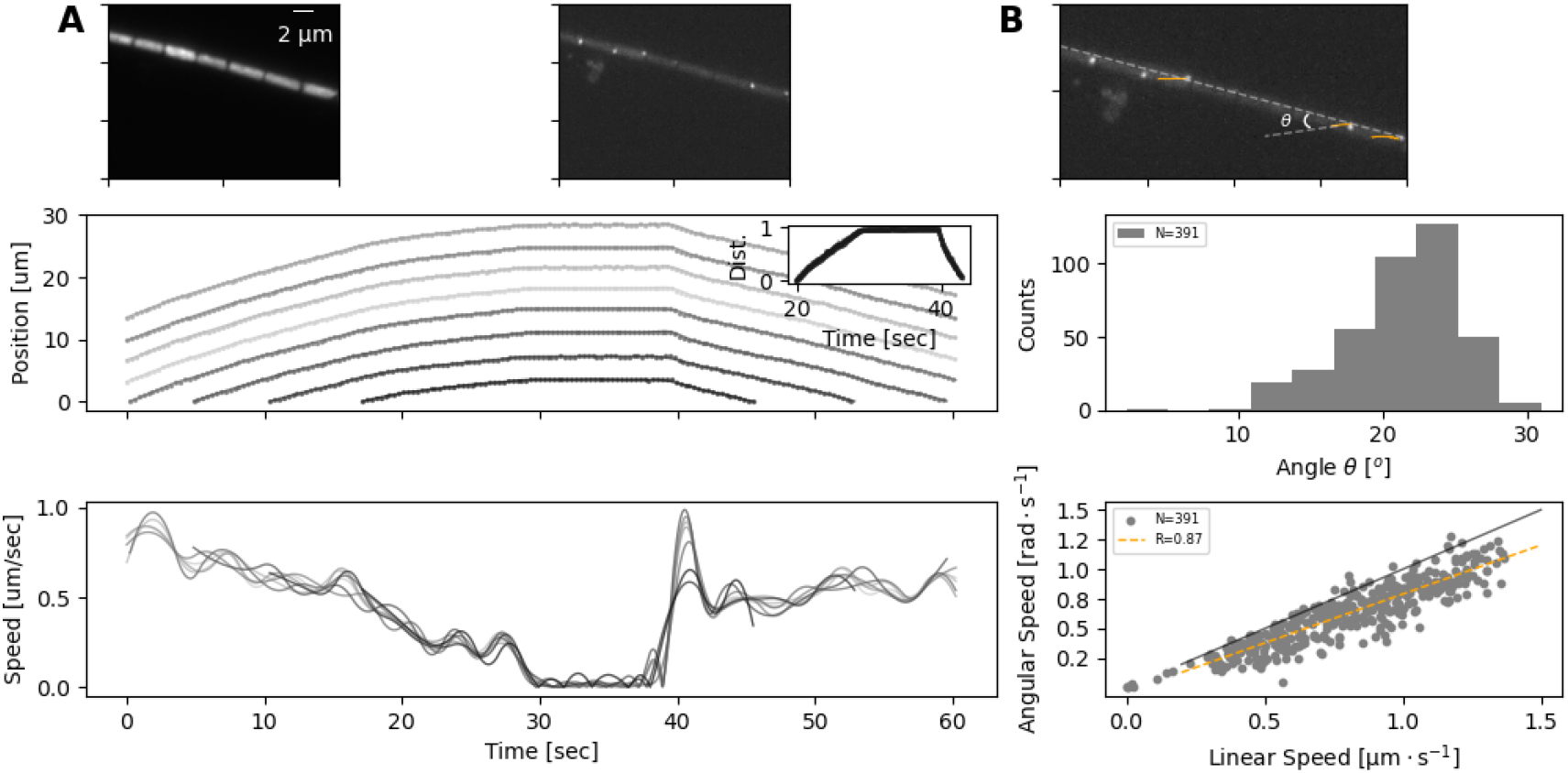
**(A) Top:** Individual images from a TIRF microscopy time-lapse, obtained with excitation using 473nm laser. The left and right panel show images with emission filters centered at 625nm and 525nm respectively. Note that these image shows a thin (≈ 200 nm) section of the cell membrane. **Bottom:** The position (top) and speed (bottom) of individual cells during a TIRF time-lapse movie (see the associated Movie S 8). Different cells’ trajectories and speed are shown in different shades of gray. The inset on the upper panel shows the normalised distance travelled by each cell, revealing high coordination in their movement.

In addition to identifying individual cells, TIRF microscopy revealed membrane-bound protein complexes localised predominantly near the poles of each cell (Fig. 4B, top panel). We have also tracked these complexes and found that their trajectories match 2-dimensional projections of a 3-dimensional helical trajectory, showing that filaments rotate as they translate (Fig. 4B, top panel and Movie S 8). The angle *θ*, that the trajectories make with the long axis of the filament shows a normal distribution with a mean of 21.2°, while the angular speed of the protein complexes are well correlated with the linear velocity of the filament (Fig. 4B). These findings indicate either a helical force generation or a screw-like structure on the cell exterior that causes a strong coupling between rotation and translation. It is interesting to note in this context, that staining filaments with fluorescently labelled Concanavalin A (see *Methods*), which binds to slime, results in a helical pattern along the filament long axis (Fig. S8). In time-lapse TIRF movies, cellular protein complexes can be seen moving through this fluorescence pattern, suggesting that the slime surrounds the filaments as a tubular structure, fixed in position relative to filament movement (Movie S 9). The tubular nature of slime is further supported by scanning electron microscopy (SEM) images, which show features around the filaments that look like collapsed slime tubes Fig. S9.

### Reversals in longer filaments can lead to buckling and plectoneme formation, causing loss of cellular coordination

The above findings show that filaments display a high level of cell-to-cell coordination, supporting the model result that cellular coordination is one of the requirements for rapid reversals. **(B) Top:** The same TIRF image as shown on the right side of top panel A, but displaying the trajectories of some of the membrane bound protein complexes (orange lines) and the angle of this trajectory with the long axis of the filament; *θ*. **Bottom:** The distribution of the angle *θ* (top) and the relation between rotational speed of the protein complexes and the linear speed of the filament (bottom). The black and orange lines in the bottom panel show the diagonal and the correlation fit (see inset) between linear and angular speed. Both angle and speed measurements are collected from 11 filaments and 391 trajectories.

In addition, we find a strong coupling between rotation and translation. Thus, it is possible that any differences in cell movement, for example the leading cells initiating reversal earlier, could lead to torsional forces developing along the filament. Supporting this, we note that cellular speeds display a steep acceleration immediately after a reversal (Fig. 4A), a characteristic observed in all filaments analysed with TIRF. In a few filaments, we have also observed indication of leading cells initiating reversal earlier than trailing ones (Fig. S10 and Movie S10). To further explore this possibility of de-coordinated movement dynamics along the filament, and emergence of associated torsional forces, we sought to analyse longer filaments with the expectation that such dynamics are expected to arise more readily in longer filaments. We found that longer filaments, where observed, readily display buckling, and sometimes these buckled loop regions twist upon themselves and form plectonemes. Almost all of the 61 observed cases of buckling and plectoneme formation involved filaments longer than 400 µm (Fig. S11).

Given that plectonemes can form, we next wanted to understand cell-to-cell coordination during such events. While it was not possible to capture a filament with TIRF microscopy, during a plectoneme formation event, we were able to record whole-filament movies at low magnification. A representative case is shown on Figure 5 (and associated Movie S 11), where we were able to analyse movement dynamics for each end of the filament. We found that the filament initially moves in a coordinated manner, on a well-defined trajectory and the two ends of the filament being coordinated in speed (Fig. 5B). With the start of buckling and plectoneme formation, however, this coordination is disrupted, with one end of the filament moving faster than the other, and finally reversing independently of the other end (Fig. 5B). This de-coordination seems to resolve towards the end of the recorded movement sequence, when the plectoneme resolves (Fig. 5 and Movie S 11). This representative case exemplifies the fact that buckling, and more specifically plectoneme formation, can result in de-coordination of cells within a filament and independent movement of filament ends. We have observed more extreme cases of such dynamics on glass, where plectonemes result in complete de-coordination in filament movement (Movie S 12). We also observed that plectonemes can get “dragged” and sometimes entangle other filaments (Movie S 13), a dynamic process that could be relevant for initiating aggregation of multiple filaments.

**Figure 5:**
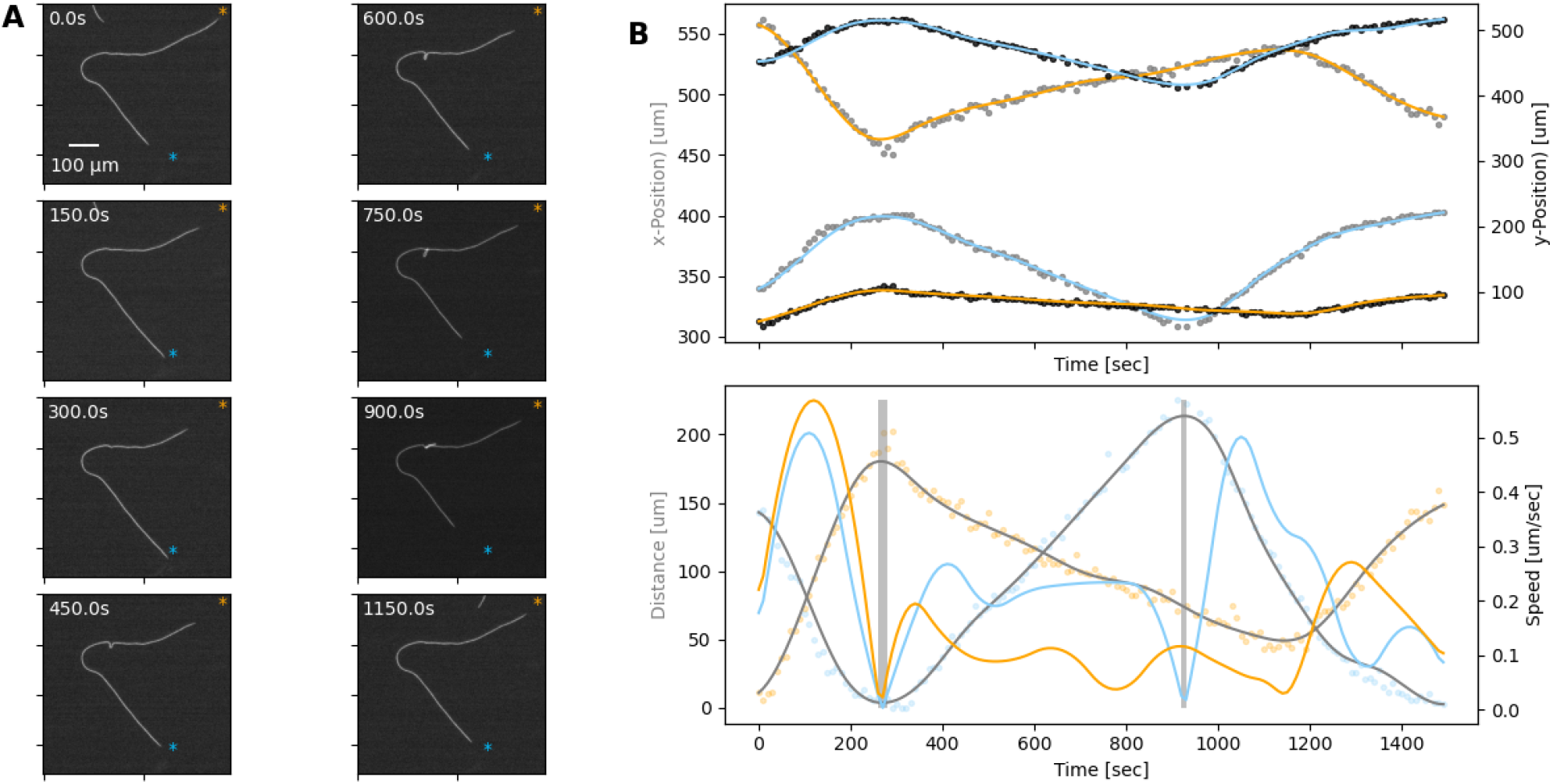
**(A)** A filament forming a plectoneme during movement under agar, shown at different time points as indicated on each panel. The scale-bar shown on the first image applies to all subsequent ones. The orange and blue asterisks indicate the end-points of the filament’s trajectory. See also the corresponding supplementary Movie S 11. **(B) Top:** The x- and y-coordinates of the two ends of the filament over time. Data points for the x- and y-coordinates are shown in gray and black respectively. Lines are spline fits to the data and their orange and blue colors indicate the filament end that stays close to the corresponding trajectory end-point shown on the images in panel A. **Bottom:** Speed and distance of each end, respective to the end-point of the trajectory close to them. Speed is shown as blue and orange lines, calculated from the respective position data shown in the top panel. Distance is shown in orange and blue data points, indicating the filament end that stays close to the corresponding trajectory end-point shown on the images in panel A, while gray lines show spline fits. The shaded areas indicate the reversals. Note that the second reversal involves only the end that stays close to the blue asterisk on panel A.

## 5 Discussion and Conclusions

Here, we analysed the motion dynamics of the filamentous, gliding cyanobacterium *F. draycotensis*. We show that gliding involves movement of filaments on curved or straight trajectories, interspersed with re-orientation or reversal events. Filament speed decreases and then increases around a reversal event, with mean speed around one micron per second, independent of filament length. We find that the time spent in reversal displays a long-tailed distribution, with most reversals taking few seconds, but some involving considerable time of filaments being stationary. We show that these experimental observations can be recapitulated by a physical model that assumes individual force generation by each cell, mechanical-coupling among cells, and mechano-sensory (coordinated) control of force direction. This model suggests that such coordination of propulsion direction among cells in a filament is important for being able to achieve fast reversals. In line with this insight, we find that cells in a filament remain highly coordinated during reversals and that their translational movement is coupled with rotation along the long axis of the filament. Following from the latter observation, we find that longer filaments can readily buckle, and buckled regions can twist upon themselves to form plectonemes. The formation of plectonemes results in de-coordination of cellular movement and independent dynamics for filament ends. We hypothesize such dynamics to be linked with the loss of sensory coupling among cells, and de-coordination of the direction of their propulsive forces. Taken together, these results provide a useful characterisation of gliding motility, which will inform future studies on both molecular mechanisms of force generation and collective behaviours of filaments.

The presented model focussed on capturing filament reversals and cell coordination during these, for which a 1D representation and mechanical coupling was sufficient. The mechanical coupling, and in particular sensing of external force fields implemented in this model, imply presence of mechano-sensing. To this end, several cyanobacterial genomes are shown to harbour homologues of the *E. coli* mechano-sensory ion channels MscS and MscL [37] and we have identified similar MscS protein sequences in *F. draycotensis*. Furthermore, *F. draycotensis*, as with other filamentous cyanobacteria, have genes associated with the type IV pili, which are implicated in the surface-based motility of filamenteous cyanobacteria [4] and in mechano-sensing in other species [38, 39]. All of these factors suggest the possibility of mechano-sensing in these cyanobacteria.

In terms of future model development, we would highlight the need for expanding into 2D or 3D models to better represent rotational and elastic bending forces and plectoneme formation. Lacking these features, the current model was still able to capture reversal events and dwell times. More detailed models, however, might be needed to fully capture reversal frequency, which we found to be independent of filament length on glass, where filament motion is not constrained. The presented, 1D model requires a dependence of cellular memory on filament length to reproduce this observation (Fig. S14-S16), which could be possible depending on molecular mechanisms of motility.

In terms of molecular mechanisms of motility and propulsive force generation, our findings that rotational and translational movement are closely coupled in *F. draycotensis* and that slime presents itself as a helical tubular structure around filaments are highly relevant. Rotation, as well as its absence, and surface fibrillar arrays have been commented on in several species of gliding, filamenteous cyanobacteria, but the rotation-translation coupling was not quantified before [27, 30, 33]. Cellular rotation has also been highlighted as a key dynamical feature in the propulsive force generation in gliding, single-celled bacteria [24, 18]. Taken together, these observations point to the possibility that rotation-translation coupling and propulsive force generation in filamentous cyanobacteria involves helical tracks surrounding the cell and either motor complexes moving on them, as suggested for single-celled, gliding bacteria [24, 18] or pili being pushed across them. Recent studies from *Nostoc* species indicate propulsion to result from pili extension [6, 32], but whether that species presents filament rotation or fibrillar arrays is unclear [30].

Photoresponse in the motility behaviour is shown in several gliding filamentous cyanobacteria [7, 8, 9, 10], but a fully quantitative analysis of these at the single filament level is currently lacking. In this context, an interesting observation we made in this study is the presence of fluorescent, membrane-bound protein complexes. We hypothesize that these complexes are associated with light sensing, since our tracking of them in TIRF microscopy always coincided with reversal of filaments. Thus, it is possible that the wavelength at which they are excited (437nm laser used in TIRF microscopy) is also acting as a sensory signal for reversals. This possibility will be explored in a future study, along with further characterization of any phototaxis behavior in *F. draycotensis*.

Gliding motility in filamentous cyanobacteria is commonly associated with aggregate or macro-scale structure formation. These formations are relevant in the context of physiological functions of the cyanobacteria, and their associated microbial communities, as shown in the case of *F. draycotensis* [13]. Similarly, in the case of the ocean-dwelling species of the *Trichodesmium* genus, aggregate formation is linked with the biogeochemically relevant processes of iron acquisition [14]. The results presented here show that de-coordination of cellular forces within a filament can lead to formation of plectonemes, which can entangle multiple filaments together. We also note that loss of motility results in the loss of aggregates in *F. draycotensis* cultures [13]. Thus, further studies of motilty and plectoneme formation can elucidate how aggregate and macro-scale formations develop and inform their physical modelling.

Given that gliding motility is observed in phylogenetically diverse bacteria, it is possible for it to be underpinned by different molecular mechanisms and present different motion dynamics. The inter-connectivity of propulsion and slime secretion is difficult to disentangle purely by molecular approaches, and development of physical models solely based on identified proteins can be limited. In particular, drastically different physical models can be constructed from observed molecular constituents. We advocate coupling of molecular studies with detailed analysis of motion dynamics, as quantification of these can provide an additional framework to constrain and test different models, ultimately providing a more complete understanding of gliding motility and associated collective behaviors.

## 6 Methods

### Sampling and Culturing

Cultures of *F. draycotensis* were grown in medicinal flasks, in volumes of 30 mL BG11+ with added vitamin mix as described previously [13]. Cultures were kept under visible light irradiance of ≈ 30 *µ*M· photons· m^*−*2^s^*−*1^, across a repeating 12-hour light/12-hour dark cycle. Samples were obtained from cultures aged between 4 and 8 weeks, by withdrawing 0.5 mL of material containing several pieces of biofilm, transferring the aliquot into a 1.5 mL microfuge tube and vortexing for 1 minute to break up the material. If the mixture was not homogeneous, additional mixing was performed using a 1 mL mechanical pipette.

### Sample Preparation for Imaging

Prior to imaging, collected innoculate was diluted into fresh BG11+ media with vitamin mix to a desired dilution (typically 1:50). For microscopy performed with 1.5− 2% agarose pads [40], 3 µL of the homogenised diluted sample was added to the pad. The droplet was then left to dry for 10-15 minutes. Where stated, the agarose pad was flipped onto a 22×50 mm cover glass, sandwiching the sample between the agarose and the glass. Otherwise, the sample was left on top of the agarose and the pad placed in a cover glass bottomed dish (MatTek/CellVis) for microscopy.

As described, some samples were imaged inside a millifluidic chamber device (Ibidi *µ*-Slide III 3D Perfusion) or on glass slides, using a 1.5 × 1 cm Gene frame (Thermo Scientific) as a spacer. For glass slide samples, the Gene frame sticker was placed on the standard glass slide. 200 µL of diluted sample was added to the frame, and the chamber was sealed with a cover glass.

Regardless of sample preparation method, all prepared samples were left to settle for 30-60 minutes, being kept on a laboratory bench under room temperature and illumination.

### Imaging Conditions

Light microscopy was performed on an Olympus IX83 microscope, using a CoolLED pE-300 illumination system. Imaging was performed using a CoolSnap-HQ2, with an imaging interval of 1-2 seconds or longer. The stage is enclosed in an environmental chamber and stabilised at 25^*°*^C. Phase imaging was performed with a low exposure rate (≈ 5 ms). Chlorophyll fluorescence was measured through a Chroma DSRed filter cube, with a recorded excitation light power of 17.4 mW and an exposure time of 100 ms. TIRF imaging was performed on a standard NanoImager system (ONI) using 473nm excitation laser and emission filters centered at 525nm and 625nm.

### Image Analysis

Images were analysed with custom-written software in Python. Analyses involved tracking individual filaments, cells, or membrane complexes on each cell, across time-lapse images. In the case of filament tracking, movies of a larger field of view were cropped to focus on single filaments, and these were processed by fitting a contour on the filament and deriving a mid-point spline from this. By repeating this process across each image in a time-lapse, it was possible to track the position of middle, trailing, and leading ends of the filament and their speed. In the case of longer time-lapse movies, any drift in the images was stabilised using stationary filaments within the larger field of view. All code used in these analyses are made available on the Github repository; https://github.com/OSS-Lab/Cyano Gliding ImageAnalysis and Model.

### Biophysical model

The model code is available at the Github repository;github.com/OSS-Lab/Cyano Gliding ImageAnalysis and Model. We model cyanobacteria as out-of-equilibrium filaments. For simplicity, we consider only a 1-dimensional model, but extension to higher dimensions are possible. The filament is composed by *N*_*f*_ beads that represent cells in a cyanobacteria filament. The approach to model the filament as connected cells with springs between them is similar to several other physical models of active filaments and polymers[41, 42, 43]. The position of bead *i* is denoted by *x*_*i*_ (*i* = 1..*N*_*f*_). Beads are connected with their nearest neighbours, along the filament, by harmonic springs. Each bead follows an over-damped equation of motion, given by;

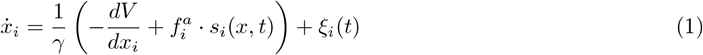

where *γ* is the friction coefficient and *ξ*_*i*_(*t*) is a Gaussian white noise, following ⟨*ξ*_*i*_(*t*)*ξ*_*j*_(*t*^*′*^)⟩ = 2*k*_*B*_*T/γδ*_*ij*_*δ*(*t*− *t*^*′*^), *T* being the temperature. The potential energy *V* (*x*) reads

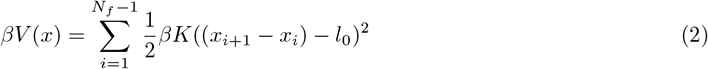

where *β* = 1*/k*_*B*_*T* is the inverse of the thermal energy (*k*_*B*_ is the Boltzmann constant) and *K* is the spring constant, representing the elastic response of the cell upon compression or extension. The rest length *l*_0_ =1 represents the unperturbed (average) length of a cell.

The remaining term in Eq. (1) represent the beads’ self-propulsion, i.e. there is a force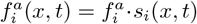, generated by the bead itself, that causes it to move in the positive or negative direction. The 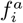 is defined as a constant in time, while *s*_*i*_(*x, t*) sets the “active state” of bead *i* at position *x* and time *t*. To model cellular direction change, we introduce a function, *ω*_*i*_(*x, t*), describing the internal “state” of each cell and dictating its active state by the relation *s*_*i*_(*x, t*) = tanh(*ω*_*i*_(*x, t*)). Notice that *s*_*i*_(*x, t*) is a continuous variable, such that −1 *< s*_*i*_(*x, t*) *<* 1. Thus, a cell can be propelled in the positive (*s*_*i*_(*x, t*) ≈ +1) or negative (*s*_*i*_(*x, t*) ≈ −1) *x* direction or it can be inert (*s*_*i*_(*x, t*) ≈0). From a biological perspective, *ω*_*i*_(*x, t*) abstracts the behavior of a cellular signalling network that integrates external and internal inputs for the cell and sets its propulsive direction. This idea is motivated by the fact that all studied uni-cellular gliding bacteria incorporate directional change, and individual cells in filamentous cyanobacteria are reported to change the location of their Type IV pili apparatus across the cell poles upon reversals [32, 6].

We define *ω*_*i*_(*x, t*) as integrating several inputs, as follows:

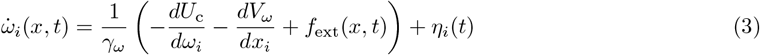

where *γ*_*ω*_ is a friction coefficient, *η*_*i*_(*t*) is a Gaussian white noise, *U*_c_ is a confinement potential, *f*_ext_(*x, t*) is an (adimensional) external force field, and *V*_*ω*_ is a mechano-sensing (adimensional) potential. The confinement potential is a mathematical abstraction that allows us to maintain the output of *ω*_*i*_(*x, t*) in a finite interval. This potential is given by:

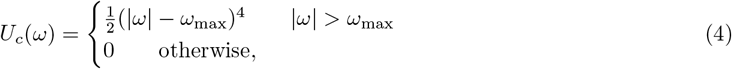

where the parameter *ω*_max_ determines the extreme values that *ω*_*i*_(*x, t*) can attain. As such, this parameter acts as a memory term, controlling a refractory period where cells would not switch direction; when given terms in Eq. (3) push a cell in one direction, *ω*_max_ will determine the time when their effects will start changing *ω*_*i*_(*x, t*). The noise term captures any fluctuations, either internal to the cellular signalling networks or caused by external noise (thermal or otherwise). It ensures that the individual cells can also turn on or off or switch direction spontaneously. The noise term *η*_*i*_(*t*) follows ⟨*η*_*i*_(*t*)*η*_*j*_(*t*^*′*^)⟩ = 2*D*_*ω*_*δ*_*ij*_*δ*(*t* − *t*^*′*^), with *D*_*ω*_ = *k*_*B*_*T*_*ω*_*/γ*_*ω*_ a diffusion coefficient, and *T*_*ω*_ being the temperature that characterises the magnitude of the fluctuations of *ω* in absence of external stimuli.

Following the experimental observations, we postulate the existence of an external, mechanical stimulus that keeps filaments on a well-defined trajectory on agar, i.e., a track. For simplicity, we assume that the track is fixed and extends from *x/l*_0_ = 0 to *x/l*_0_ = *L*. The external stimulus field is given by the following functional form:

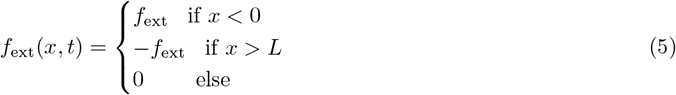

where *f*_ext_ is an adimensional constant. Given Eq. (3), the action of the external field drives the auxiliary variable to positive/negative values. When considering filaments on glass, we simply set this external stimulus to zero.

The potential *V*_*ω*_ encodes the mechano-sensitivity of each cell to their neighbors and is given as;

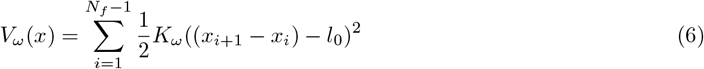

where the quantity *K*_*ω*_ = *f*_ext_*/l*_0_ + *δK*_*ω*_ is an effective spring constant. The function *V*_*ω*_ encodes the feedback to a mechanical stress (compression / extension) applied to each cell, compelling the direction of self-propulsion so that this stress is relieved.

Notice that, within the model, *f*_ext_ essentially encodes the basal strength of the response of the cells to stimuli, either extra- or inter-cellular. The mechano-sensing can be more (*δK*_*ω*_ *>* 0) or less (*δK*_*ω*_ *<* 0) relevant than the external stimulus; naturally, we impose *δK*_*ω*_ *>*−| *f*_ext_|, i.e., there can not be a positive feedback to compression.

### Simulation details

We integrate the equations of motion Eq. (1) using a stochastic predictor-corrector algorithm, while the equations Eq. (3) are integrated using the Euler-Maruyama algorithm. In both cases, we employ an integration time step 10^*−*3^*τ* ≤Δ*t* ≤5 ·10^*−*3^*τ, τ* being the unit of time. As mentioned above, we set the average length of an unperturbed cell as the unit of length and set the unit of time as the diffusion time of a single inactive cell 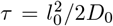 _0_. For convenience, we set the unit of energy as the active “work” *ϵ* = *f* ^*a*^*l*_0_ = 1; as such, we fix *f* ^*a*^ =1 throughout the simulations. We further fix the friction coefficient to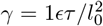. We assume that the motion of the filament is dominated by its self-propulsion, i.e. *f* ^*a*^*l*_0_ ≫ *k*_*b*_*T* and we set *k*_*B*_*T* = 0.01*ϵ*.

Concerning the parameters that enter in the evolution of *ω*, the scale of the “energy” should be set here by *k*_*B*_*T*_*ω*_. However, we set *k*_*B*_*T*_*ω*_ = 0.1 for convenience; we find that reasonably smooth reversals can be obtained for such a choice, maintaining the other parameters within the same order of magnitude. As such, we fix *f*_ext_ = 10.

### Movement characterization

The analysis of experimental and simulated trajectories focuses on the dwell time and the number of reversals. The dwell time is computed as follows: the trajectory is smoothed via a spline and the velocity is computed via numerical differentiation. A dwell time (or dwell event) is defined as the interval within which the velocity remains, in absolute value, below 15% of its maximal value. A reversal corresponds to a change of sign of the centre of mass velocity of the filament after dwelling; we name *n*_*r*_ the number of reversals counted along a trajectory. Similar to re-orientations in experiments on glass, we also identify stop-continue events when removing the boundaries: here, the centre of mass velocity does not change sign after dwelling. We define two quantities: the reversal efficiency *M* and the reversal rate *ν*. The reversal efficiency aims to capture the relation between the “ideal” and observed number of reversals, hence a value of one indicates expected behavior of a filament on a defined track. Mathematically, it is defined as;

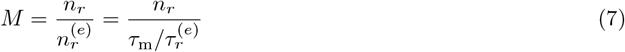

with 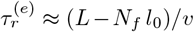 and *τ*_m_ the measurement time (either in experiments or simulations). The quantity 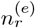 is the number of reversals one would expect from an ideal filament, that reverses smoothly and deterministically in a negligible time and then travels at constant speed. In simulations, *v* = *f* ^*a*^*/γ* = 1*l*_0_*/τ* while in experiments we estimate *v*≈ 1 µm s^*−*1^. The quantity 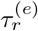 is, thus, the ideal time between two reversals, i.e. the time the centre of mass would take to travel from its initial condition (after a reversal) to the track boundary. By definition, this quantity pertains only to filaments confined between the track’s ends. Conversely, the reversal rate can be defined also without the track boundaries: it is simply defined as the number of reversals over the measurement time

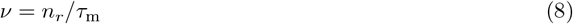

Finally, we define a syncronisation index *S* as

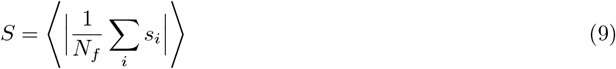

where the average is performed over time or over the different realisations. Essentially, for each conformation, we compute the (instantaneous) average propulsion direction of each cell; as we are interested in the overall synchronisation of the filament regardless of its direction, we take the absolute value.

## 8 Acknowledgements

We thank Douglas Risser and Jonasz Slomka for insightful comments on an earlier version of this manuscript. We acknowledge the help of imaging facility manager Ian Hands-Portman with microscopy setups and Nicole Robb for allowing access to her groups’ TIRF microscopy. We would also like to acknowledge the University of Warwick Electron Microscopy RTP for assistance in the research described in this paper.

## 10 Supplementary Information

### 10.1 Supplementary videos

1. **Movie S1**-Filament moving under 1.5 % agarose sandwich. Magnification of ×4 (1.61 *µ*m/pixel). Time interval of 5 seconds. Fluorescence recorded in Red channel with 100 ms exposure. AVI produced with 20 fps compression. Link: https://youtu.be/RsANG2RBzTg
2. **Movie S2**-Filaments moving across glass slide. Magnification of ×10 (0.645 *µ*m/pixel). Time interval of 10 seconds. Fluorescence recorded in Red channel with 200 ms exposure. AVI produced with 20 fps compression. Link: https://youtu.be/KOcwKboCFZg
3. **Movie S3**-Single filament in track on 2.5 % agarose. Magnification of × 10 (0.645 *µ*m/pixel). Time interval of 5 seconds. Phase image with 5 ms exposure. AVI produced with 20 fps compression. Link: https://youtu.be/XVIv4FLYBas
4. **Movie S4**-Single filament on a circular trajectory on 1.5 % agarose. Magnification of × 4 (1.61 *µ*m/pixel). Time interval of 5 seconds. Fluorescence recorded in Red channel with 100 ms exposure. AVI produced with 20 fps compression. https://youtu.be/PJdFEE6R6fk
5. **Movie S5**-Several filaments in track on 2.5 % agarose. Magnification of × 10 (0.645 *µ*m/pixel). Time interval of 5 seconds. Phase image with 5 ms exposure. AVI produced with 20 fps compression. Link: https://youtu.be/hu-U810RMuw
6. **Movie S6**-Simulation movie of an active filament with a parameter set, resulting in coordinated reversals. The filament is composed of 8 beads (cells) in a track of length 50 units. AVI produced with 5 fps compression. Link: https://youtu.be/nyUmk8QXz1I
7. **Movie S7**-Simulation movie of an active filament with a parameter set, resulting in erratic reversals. The filament is composed of 8 beads (cells) in a track of length 50 units. AVI produced with 5 fps compression. Link: https://youtu.be/Llc5oOmdxm0
8. **Movie S8**-TIRF imaging of filament undergoing a reversal. This movie is associated with the analysis shown in Fig. 4. Magnification of × 100 (0.116 *µ*m/pixel). Time interval of 33 ms. Fluorescence in cyan is from the bundled complexes on the filament surface, observed between 500-560 nm, with 30 ms exposure from a 473 nm laser. Fluorescence in the red channel comes from chlorophyll fluorescence, with 30 ms exposure from a 473 nm laser and observed in a channel between 600-670 nm. AVI produced with 100 fps compression. Link: https://youtu.be/Doa-jOkl04I
9. **Movie S9**-TIRF imaging of stacked filament in a stained slime sheath. Magnification of × 100 (0.116 *µ*m/pixel). Time interval of 33 ms. A 473 nm laser with an exposure time of 30 ms was used for excitation. The slime sheath is stained with concanavalin AlexaFluor-488 and coloured in yellow (observed 500-560 nm). Chlorophyll autofluorescence is coloured in cyano (observed 600-670 nm). AVI produced with 60 fps compression. Link: https://youtu.be/mYoqSqK9re0
10. **Movie S10**-TIRF imaging of filament undergoing a reversal, where fluorescence shows the protein complexes on the surface, This movie is associated with the analysis shown in Fig. S10. Magnification of × 100 (0.116 *µ*m/pixel). Time interval of 33 ms. Fluorescence observed between 500-560 nm, with 30 ms exposure from a 473 nm laser. AVI produced with 100 fps compression. Link: https://youtube.com/shorts/pa5iIcua8Ps?feature=share
11. **Movie S11**-Buckling filament under 1.5 % agarose. This movie is associated with the analysis in Fig. 5 of the main text. Magnification ×4 (1.61 *µ*m/pixel). Time interval of 5 seconds. Fluorescence recorded in Red channel with 100 ms exposure. AVI produced with 20 fps compression. Link: https://youtu.be/EmfIGJiiWUg
12. **Movie S12**-Filament twisted in middle, forming plectonemes on glass. Magnification of ×4 (1.61 *µ*m/pixel). Time interval of 30 seconds. Fluorescence recorded in Red channel with 100 ms exposure. AVI produced with 20 fps compression. Link: https://youtu.be/VzHMagiGqfo
13. **Movie S13**-Filaments forming plectonemes on glass. Magnification of ×4 (1.61 *µ*m/pixel). Time interval of 5 seconds. Fluorescence recorded in Red channel with 100 ms exposure. AVI produced with 20 fps compression. Link: https://youtu.be/OBm0IpUN6rI

## 10.2 Supplementary figures

**Fig. S1:**
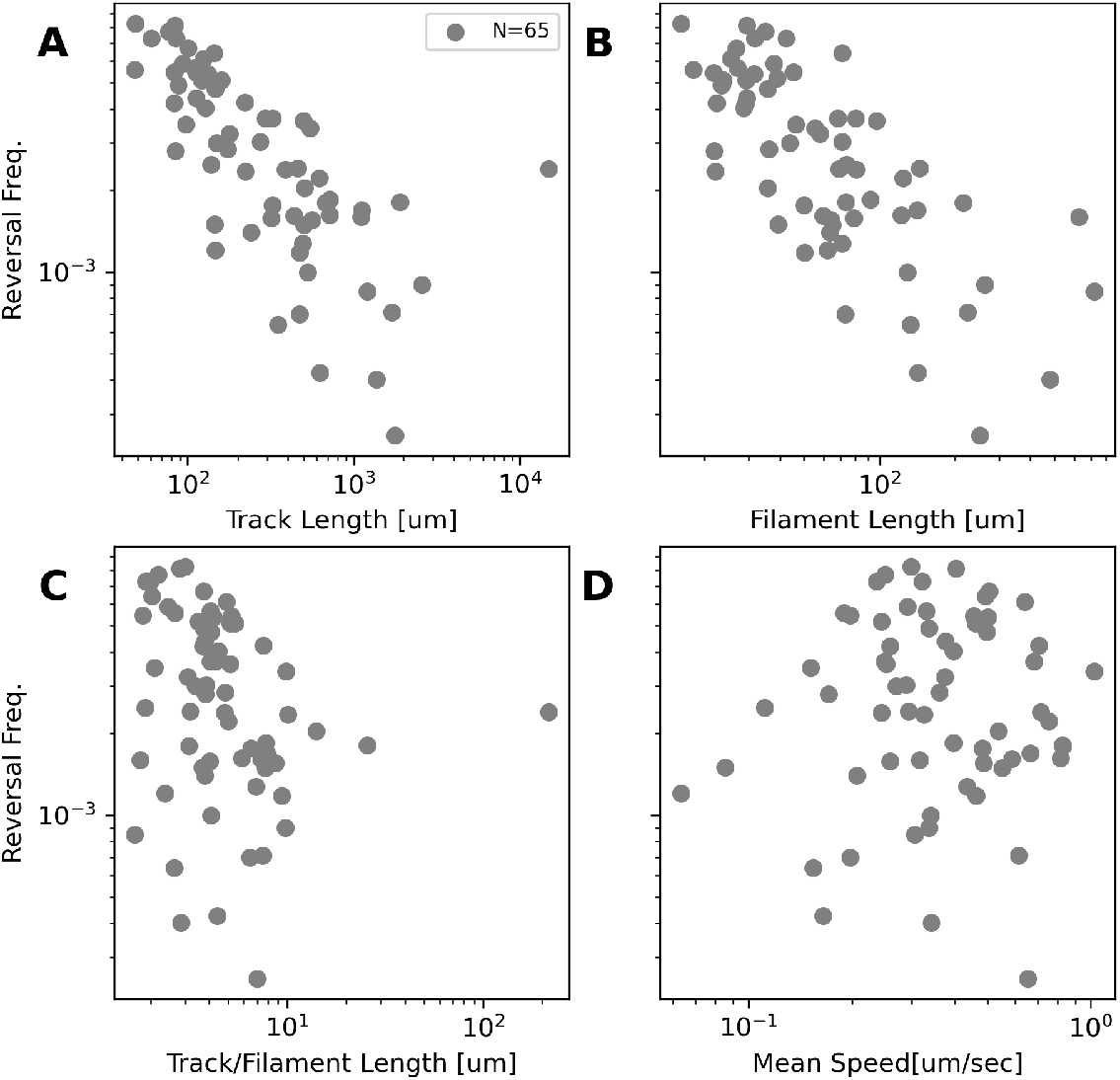
**A**. Reversal frequency against track length, determined from extreme boundaries of filament movement during observation time. Data collected from 65 filaments observed as moving under agar. **B**. Reversal frequency against filament length. **C**. Reversal frequency against filament length normalised by track length. **D**. Reversal frequency against filament mean speed.

**Fig. S2:**
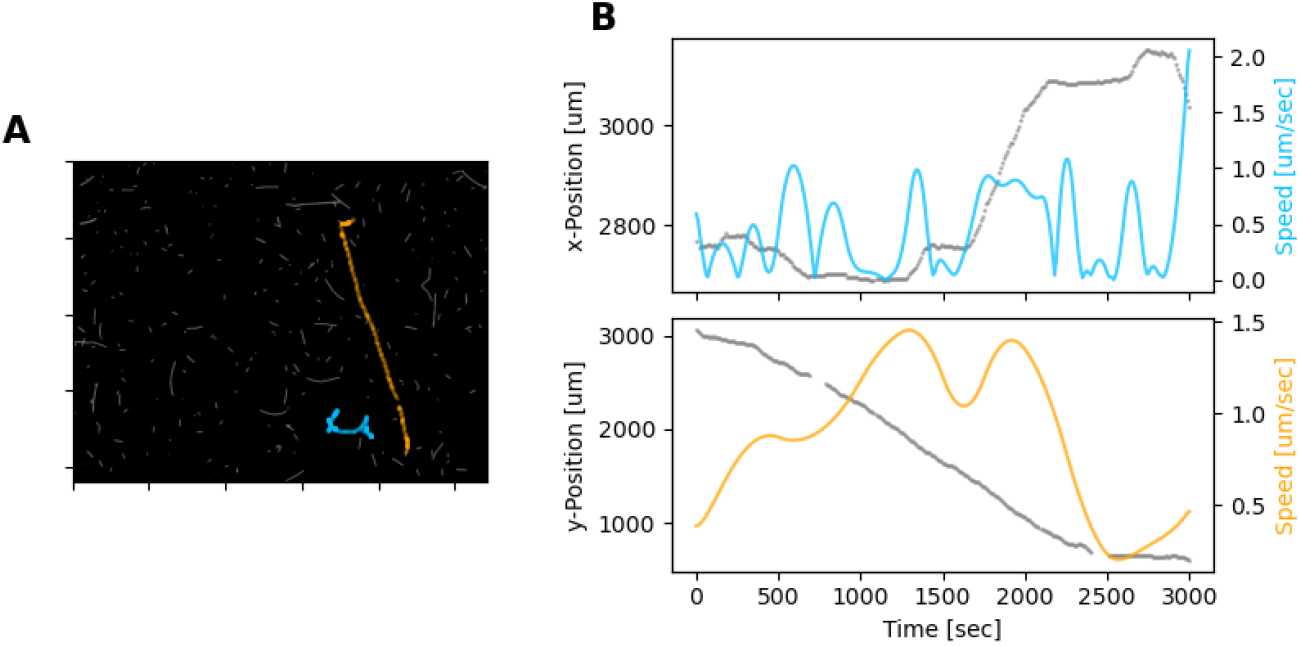
**A**. Single frame from a movie, showing filaments on glass (this image is associated with Movie S 2). Two selected filaments are highlighted with their trajectories, colored in blue and orange. **B**. Trajectory (blue and orange) and speeds (gray) of filaments shown in panel A. Trajectory colors are matching to panel A. Trajectories are only shown in terms of either the x- or y-axis for simplicity, with the axis the the most significant movement shown.

**Fig. S3:**
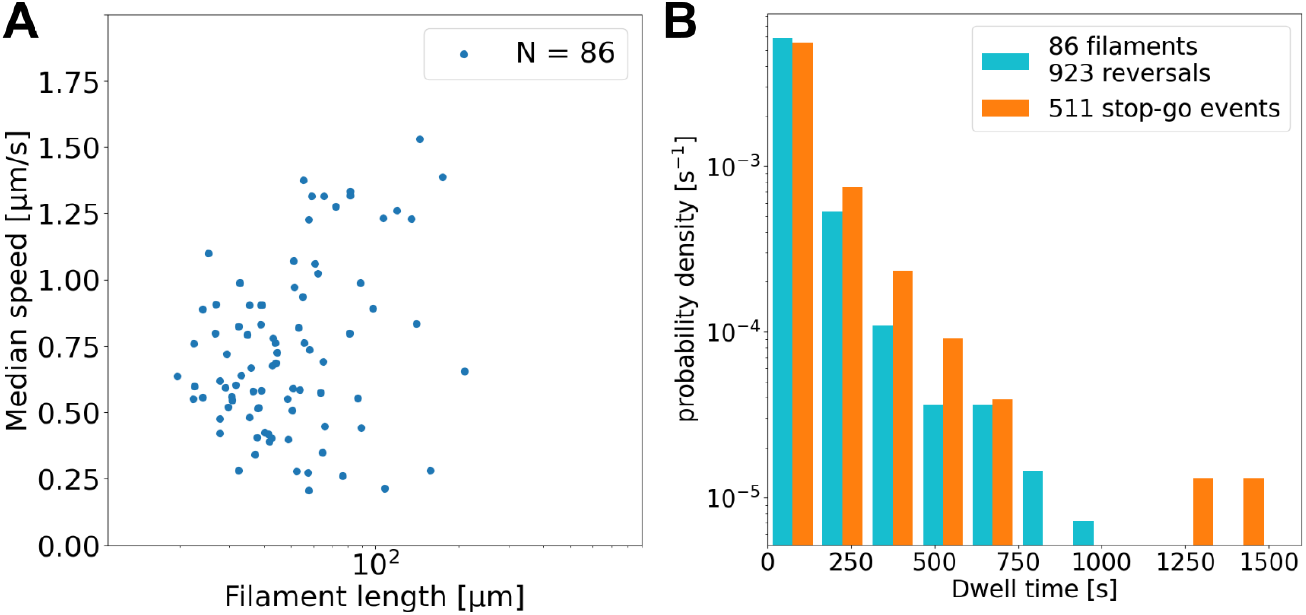
**(A)** Median speed against mean filament length for filaments moving on glass. Data from 86 individual filaments. There is a weak positive correlation between filament speed and length, which supports the conclusion that multiple (or all) cells contribute to propulsion. **(B)** Distribution of dwell times (the duration of stopping events) for movement on glass. On glass, we observe an additional behaviour following a stopping event, where the filament continues in the same direction instead of reversing, which we call a ‘stop-go’ event. The dwell time histograms for the reversals (blue) and the stop-go (orange) events are similar in shape, and there are approximately twice as many reversals as stop-go events.

**Fig. S4:**
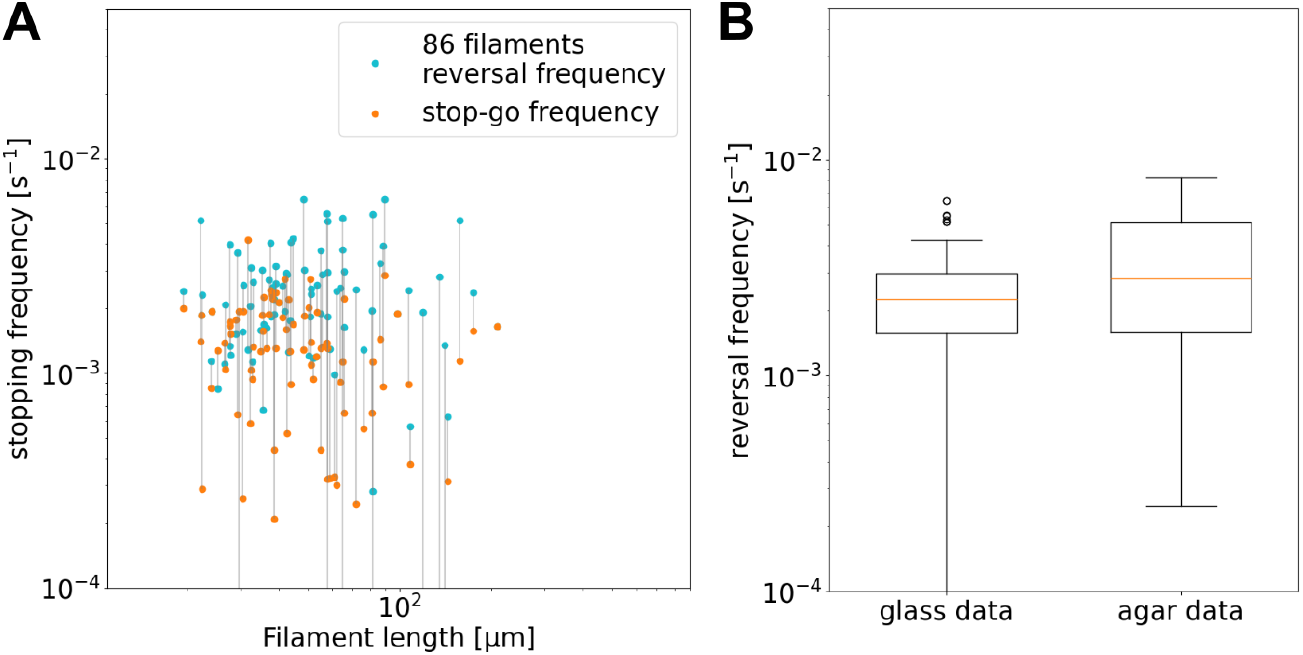
(**A**) Reversal and stop-go frequency as a function of filament length on glass, split up for each filament into ‘stop-reverse’ (blue) and ‘stop-go’ (orange) events, depending on the direction of motion following the stopping event. The two points for each individual filament are joined by grey lines (note that some of the lines are not connected, due to some filaments analysed showing either no reversals or no stop-go events, over the entire track, giving a frequency of 0 for those points, which are therefore undefined on this logarithmic axis.)The reversal frequency is generally higher than the stop-go frequency. Frequencies are approximately constant with filament length, with an average of one reversal every 410 s, in contrast to the agar data. (**B**) Distribution of reversal frequency data from observations on glass and agar. On agar, the highest reversal frequencies (mostly observations from shorter filaments/tracks) are higher than the average on glass, while the lowest frequencies (mostly due to longer filaments/tracks) are comparable to the lower glass values. This corroborates the hypothesis that on agar the reversals are caused by the track ends, so that shorter filaments are forced by the short tracks to reverse more often than their intrinsic reversal frequency (suggested by the glass data).

**Fig. S5:**
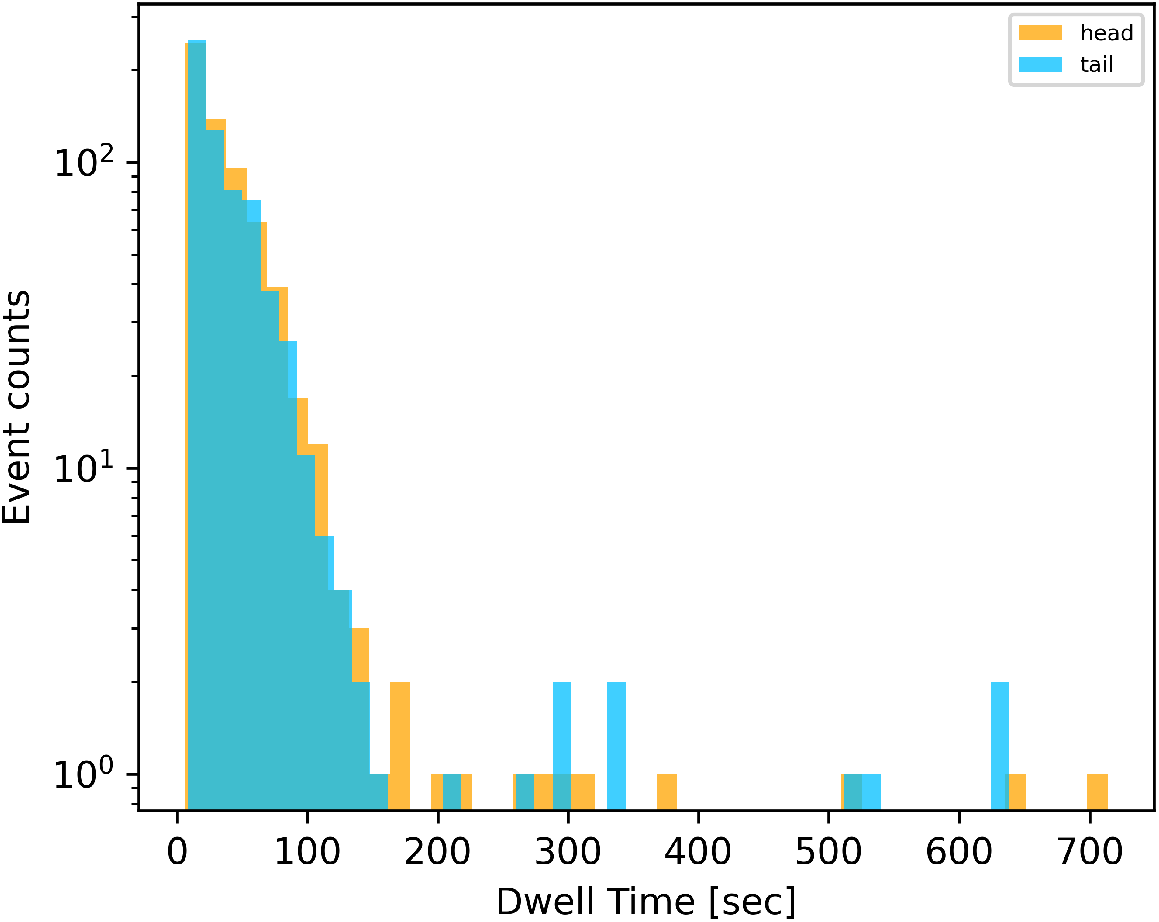
Distribution of dwell times of leading (head) and trailing (tail) ends of filaments. Data is collated from 632 reversal events across 65 filaments. Only those reversals, where dwell times estimated from tracking filament ends differed less than a threshold from those estimated from filament centre, are included.

**Fig. S6:**
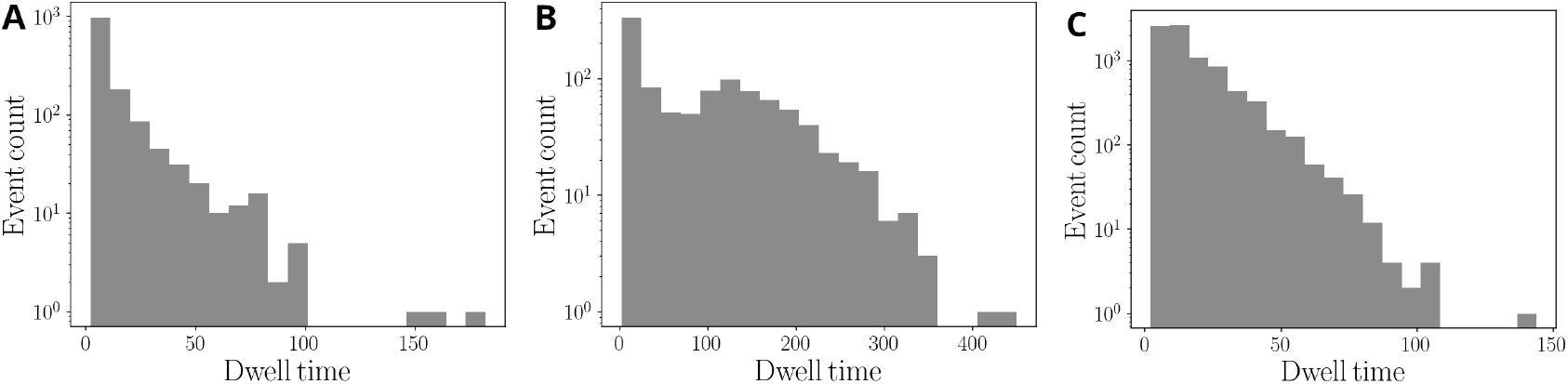
Simulated distribution of dwell times. The parameters values are as in Fig. 3C: **(A)** *K*_*ω*_ = 20, *ω*_max_ = 10; **(B)** *K*_*ω*_ = 1, *ω*_max_ = 30; **(C)** *K*_*ω*_ = 30, *ω*_max_ = 1.

**Fig. S7:**
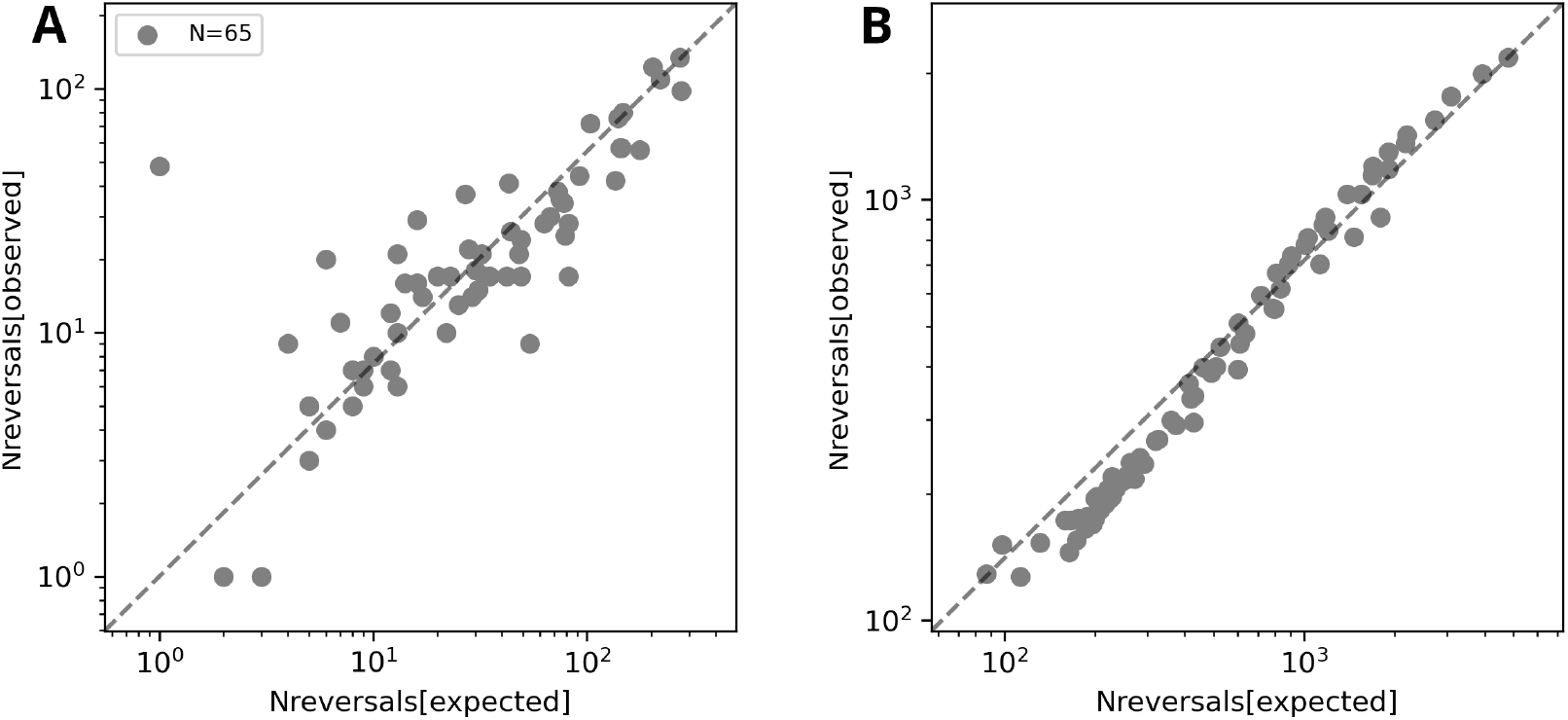
Number of reversals *n*_*r*_ against the number of expected reversals 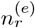 as observed in experiments under agar **(A)** or in simulations **(B)**. One outlier filament in the experimental data involved reversals prior to track ends, but in a consistent location in the track. Simulated data have been obtained with *K*_*ω*_ = 20 and *ω*_max_ = 10, 20 and several different values of *L*.

**Fig. S8:**
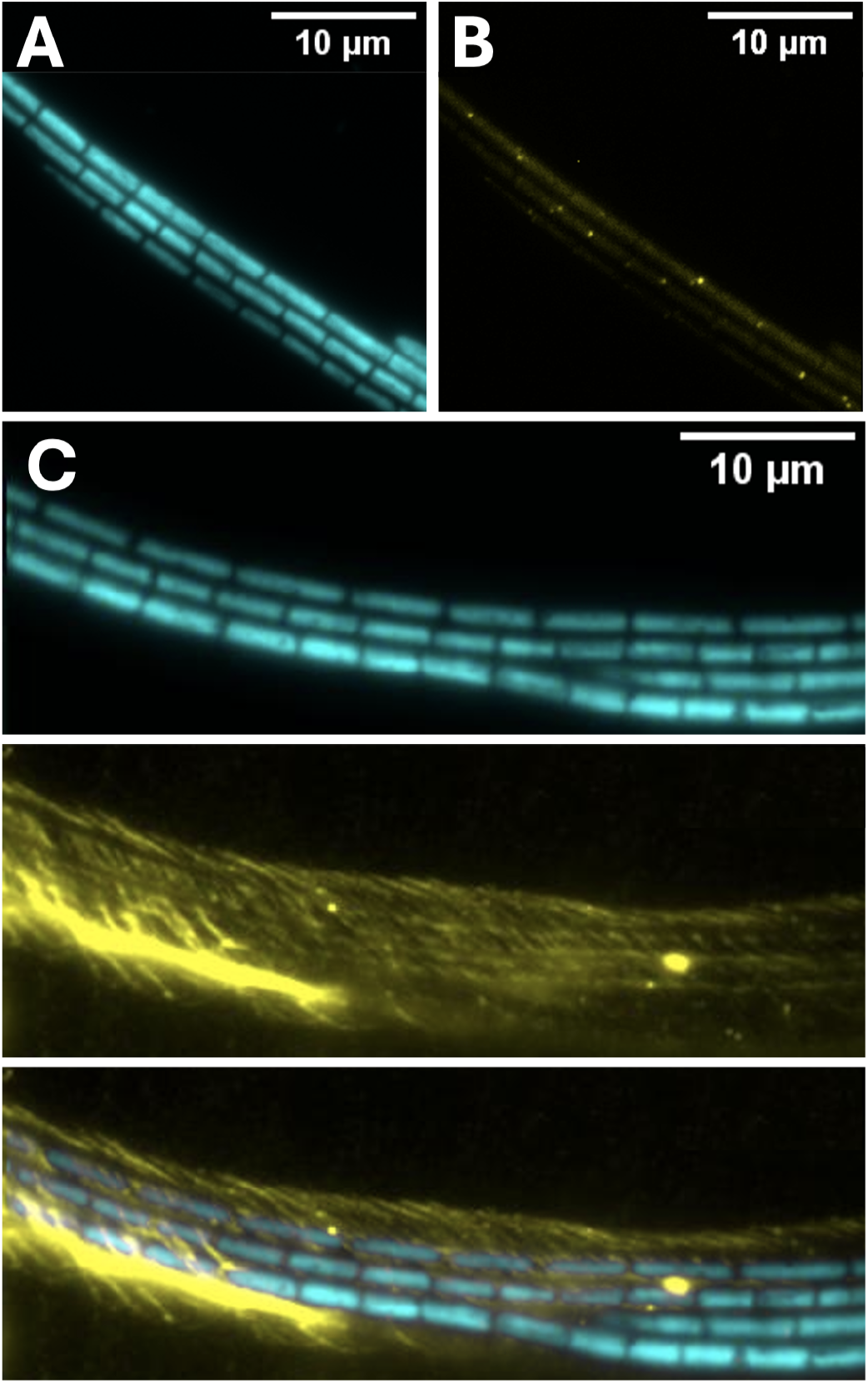
**(A)** TIRF image of *F. draycotensis* filaments illuminated with light of wavelength 473nm and imaged under an emission filter centred on 625nm. **(B)** Same filaments under an emission filter centred on 525nm, showing regularly placed fluorescent protein complexes. **(C)** TIRF image of a bundle of filaments stained using Concanavalin-A and illuminated with light of wavelength 473 nm. Top and middle panels show images with emission filters centred on 625 and 525nm, respectively. Surrounding sheath of exopolysaccharide is visible (false colored in yellow). The third panel shows the composite of these two emission channels.

**Fig. S9:**
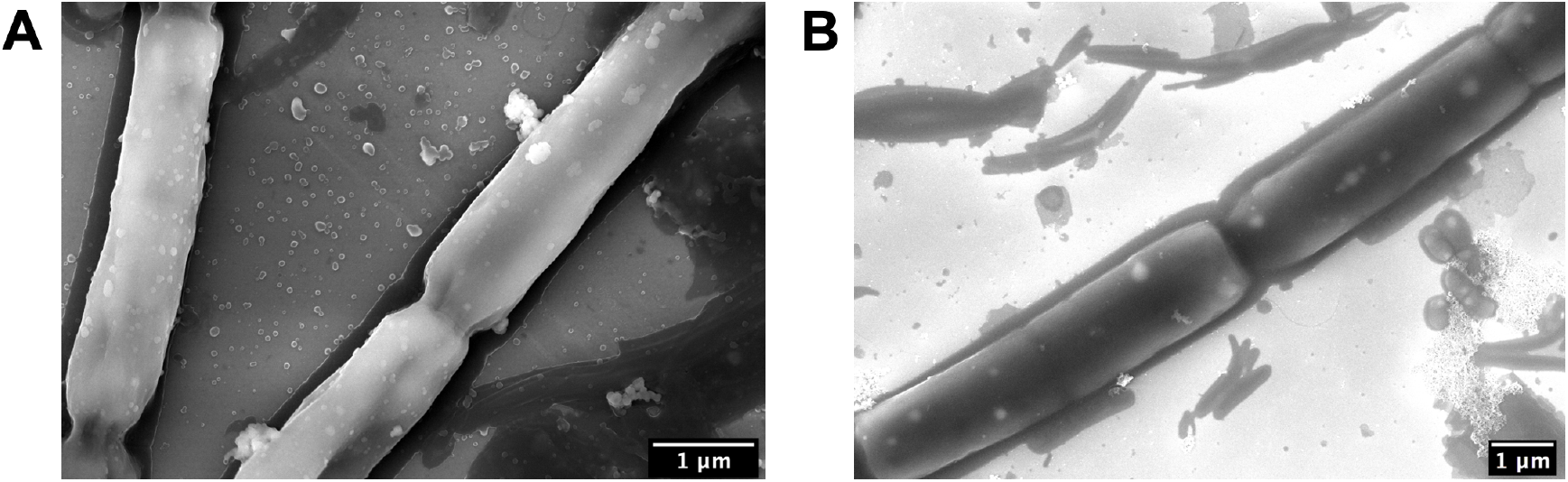
Scanning electron microscopy (SEM) images of the *F. Draycotensis* cyanobacteria filaments. (**A**) cyanobacteria filaments without further treatment, Mag.: × 40 k, EHT: 8 kV; (**B**) filaments following plasma cleaning inside SEM chamber, Mag.: × 25 k, EHT: 8kV. Performed on a Zeiss Gemini, imaged on silicon wafer

**Fig. S10:**
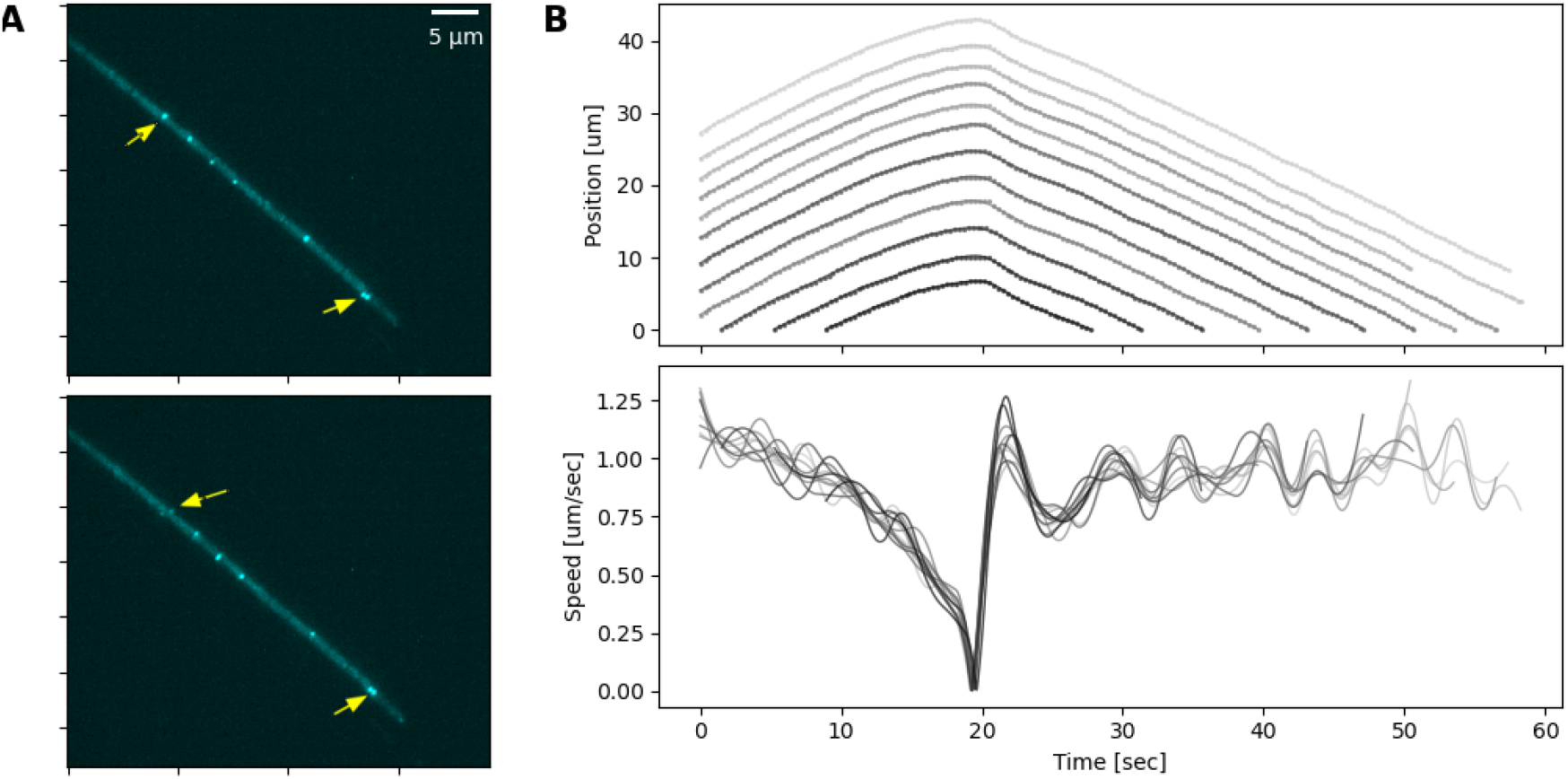
**(A)**. Two time points in the movement sequence of a filament, the position and speed data for which is shown in panel B. The location of two protein complexes are shown with arrows, indicating that the complex further up the filament starts its movement earlier than the one at the tip of the filament. See also the associated Movie S10. **(B)** The position (top) and speed (bottom) of individual cells. Different cells’ trajectories and speed are shown in different shades of gray.

**Fig. S11:**
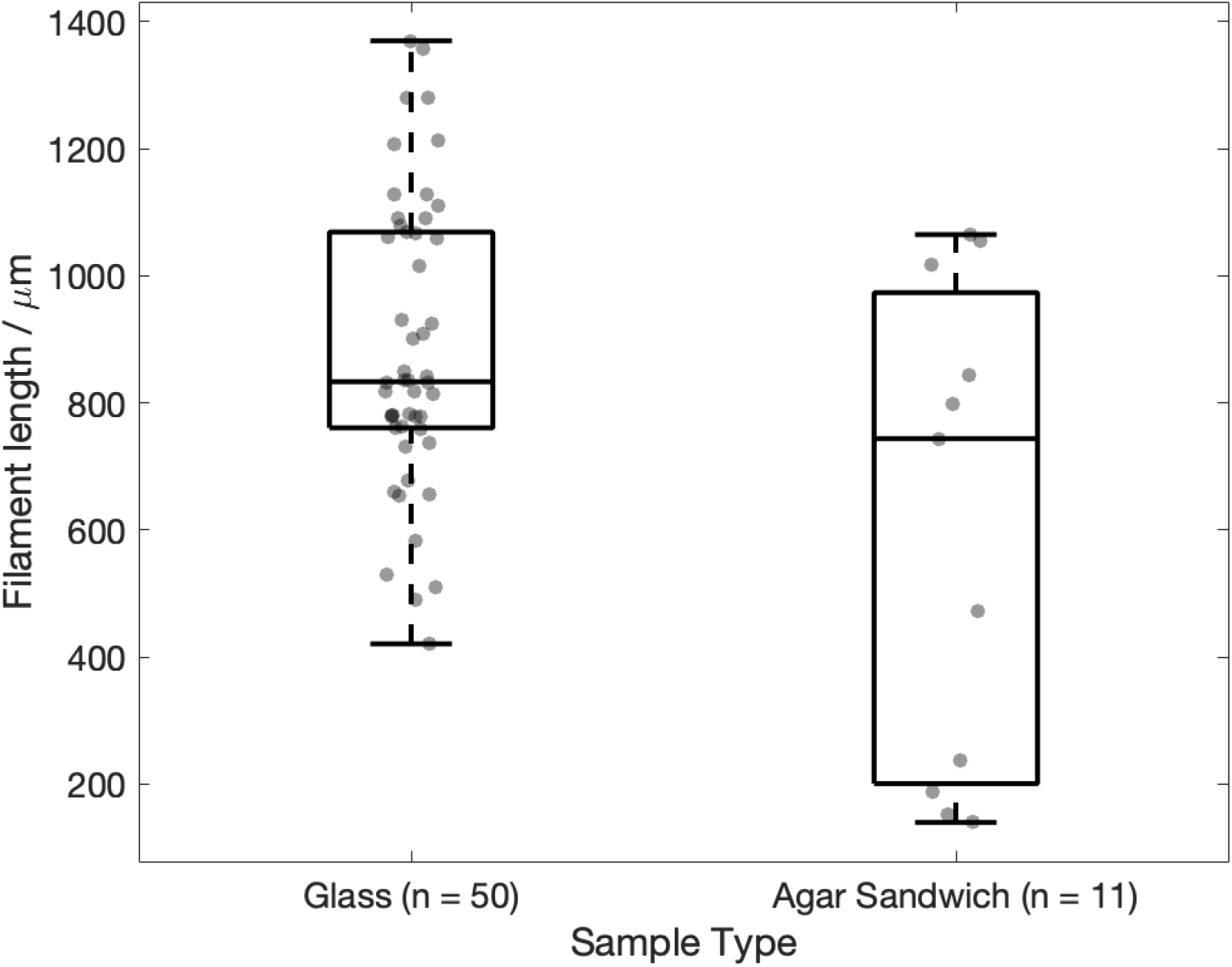
Distribution of filament lengths that exhibit buckling or plectonemes, shown for filaments observed on a glass slide (*n* = 50), or sandwiched under agar (*n* = 11)

**Fig. S12:**
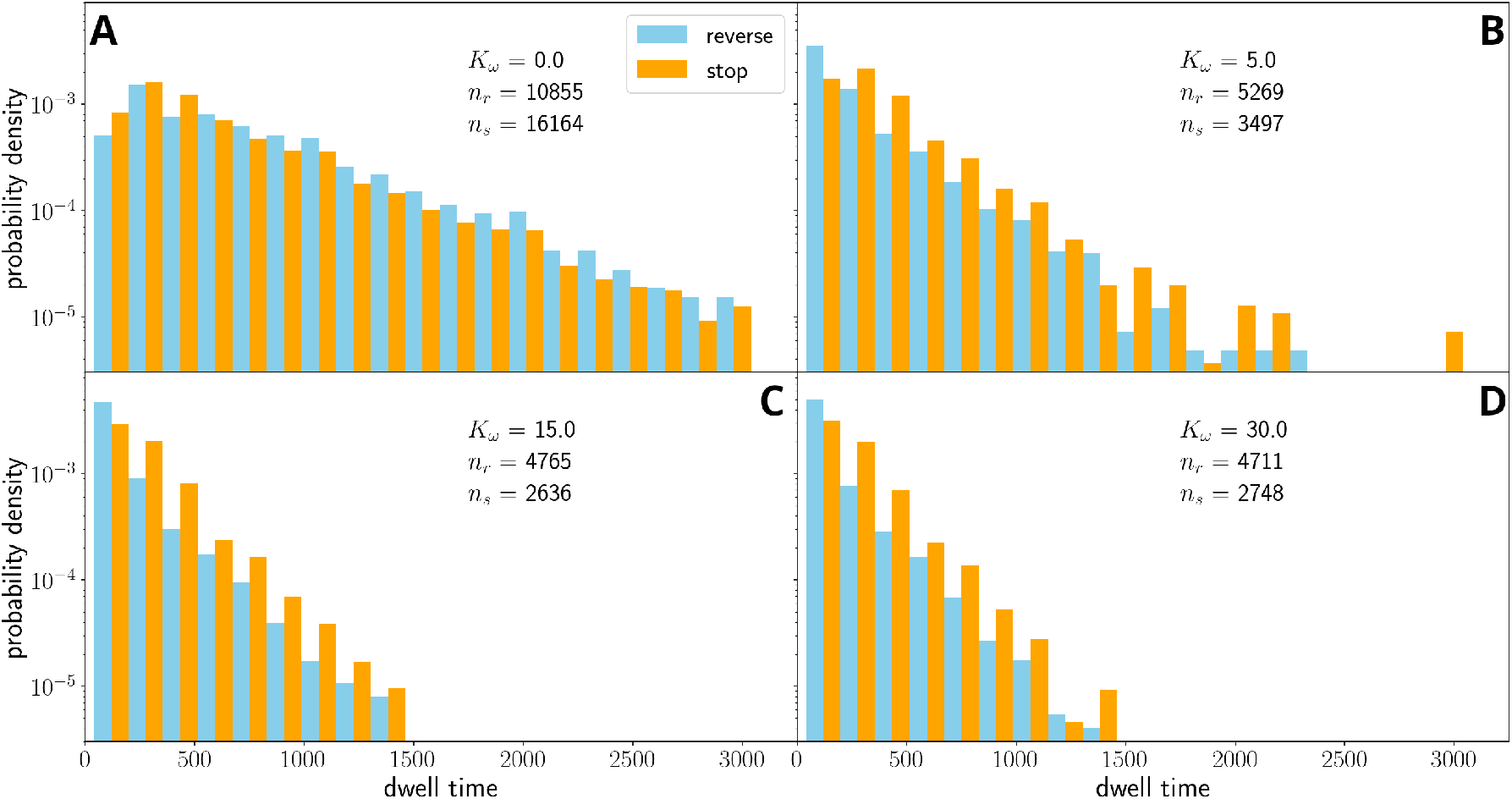
Distribution of stop/dwell times for simulated filaments on ‘glass’, i.e., without external forces. We combine data from filaments with *N*_*f*_ = 10, 50, 100, 300 (as in Fig. S3), use a constant *ω*_max_ = 10 and different values of *K*_*ω*_. Notice that, without coupling the cells’ propulsion (*K*_*ω*_=0, panel **(A)**), the distribution of dwell times develops a peak at some finite dwell time and its tail becomes significantly fatter than in experiments. More importantly, the number of stop-go events becomes larger than the stop-reverse events, which does not match the experimental evidence. With increasing coupling (panels **(B)**-**(D)**) the distribution becomes markedly exponential and the ratio between stop-go and stop-reverse instances matches the experimental value.

**Fig. S13:**
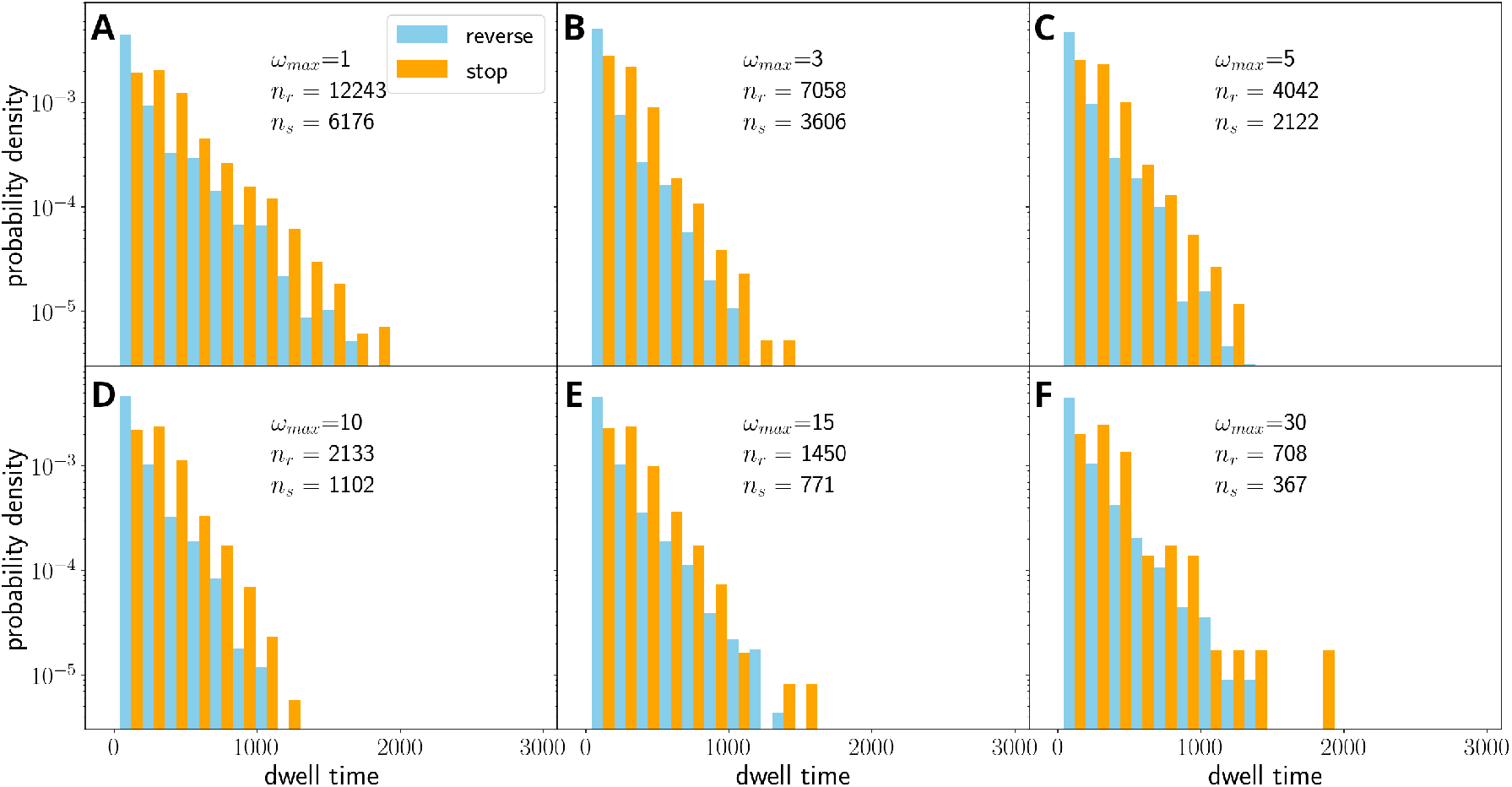
Distribution of stop/dwell times for *N*_*f*_ =100, *K*_*ω*_=25 and several values of *ω*_max_, for simulations without external force field. Above a certain threshold (panels **(B)**-**(F)**), the dwell time distribution is independent of the parameter *ω*_max_. However, the number of stops or reversals decreases, as it is harder to de-coordinate the filament.

**Fig. S14:**
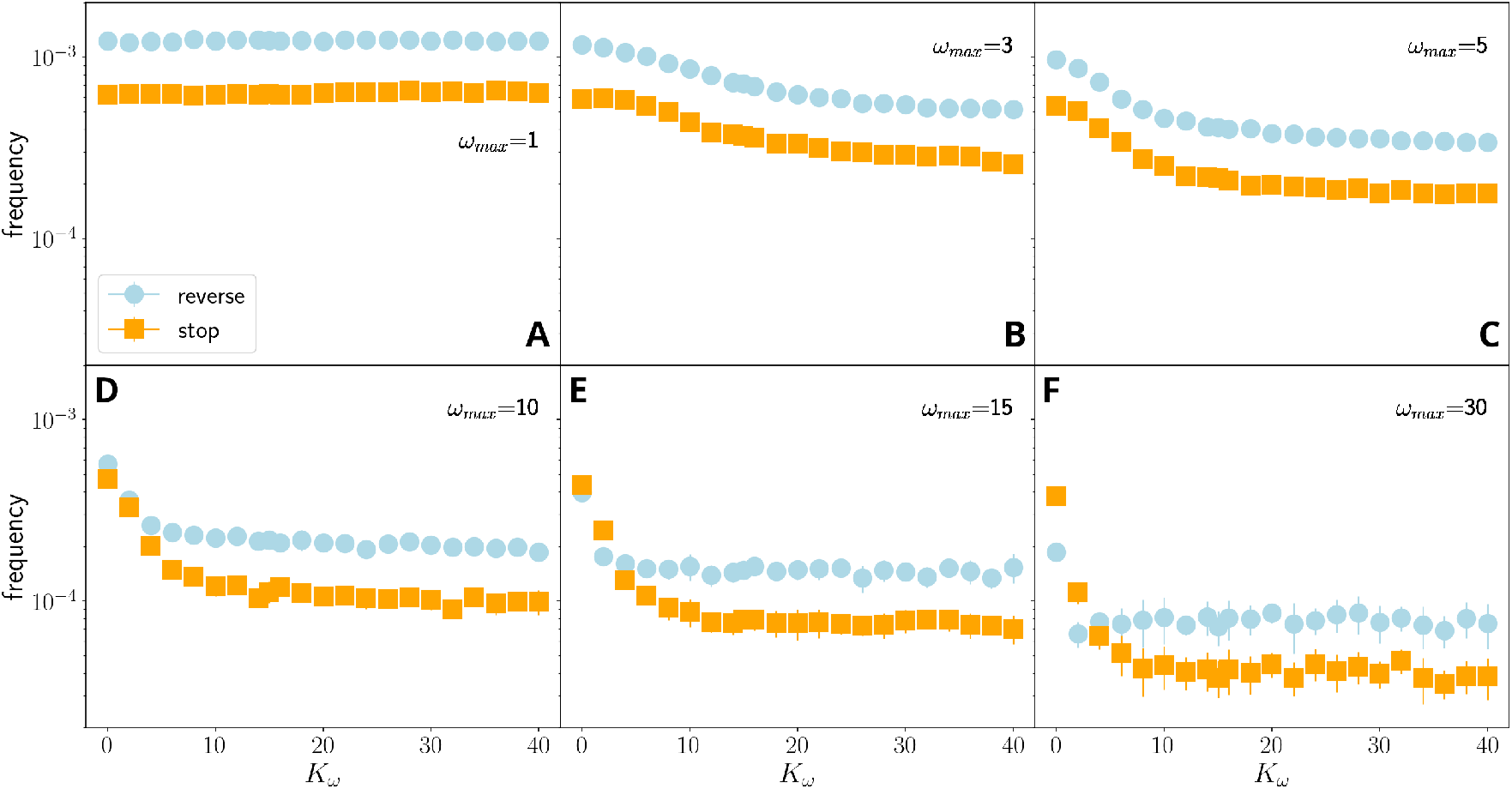
Average reversal frequency for stop-reverse (blue) and stop-go (orange) events as a function of *K*_*ω*_ for *N*_*f*_ =100 and different values of memory parameter, *ω*_max_, for simulations without external force field. At very small values of *ω*_max_ (**(A)**, negligible memory) the frequencies of stops and reversals do not depend on *K*_*ω*_, that is, the filament is not coordinated. With increasing *ω*_max_ (**(B)**-**(F)**), the frequency decreases with increasing *K*_*ω*_ and *ω*_max_.

**Fig. S15:**
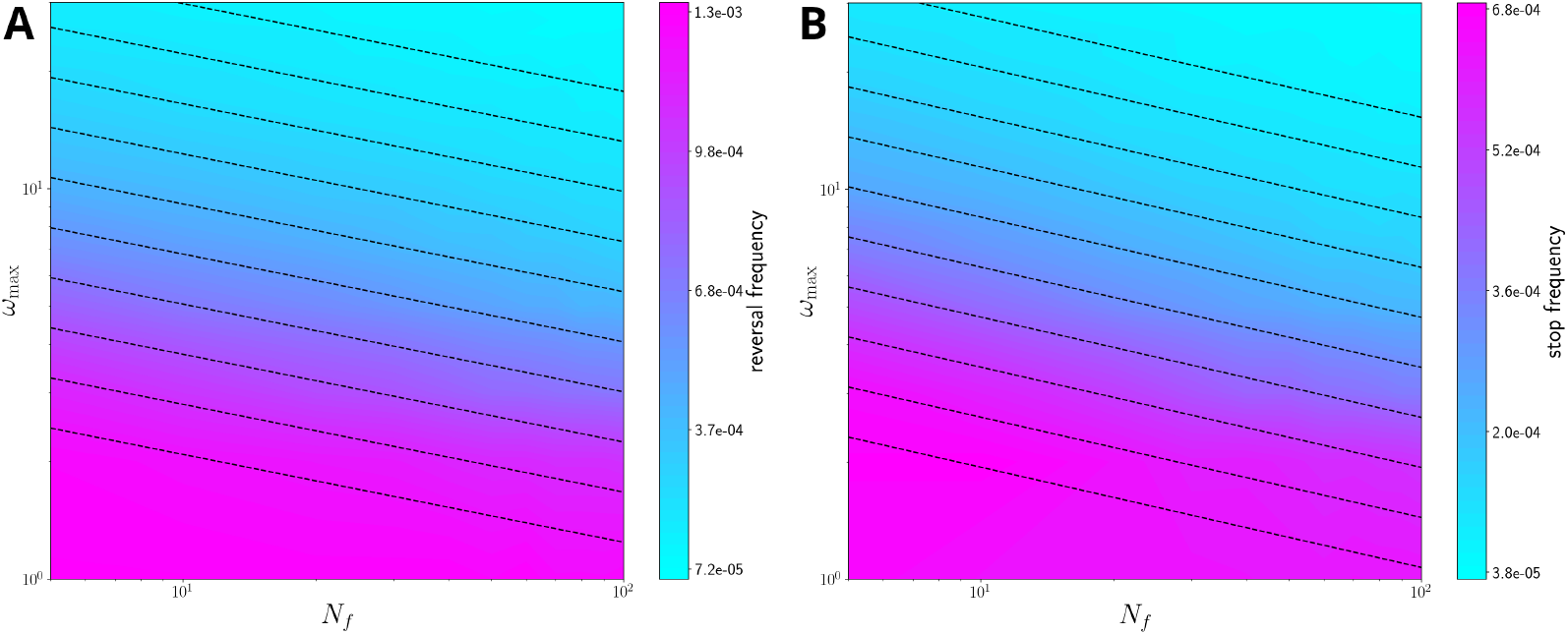
The relation of reversal frequency **(A)** and stop frequency **(B)** with filament length and memory parameter (i.e., in the *N*_*f*_− *ω*_max_ plane) at fixed *K*_*ω*_=15, for simulations without external force field. Note that the reversal frequency is always higher than the stopping frequency. The black dashed lines are approximated level curves, obtained by fitting the relationship between *ω*_max_ and *N*_*f*_ at fixed reversal/stopping frequency as a power law; in particular we obtain the phenomenological relations ∝ 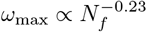 for the reversal frequency and *ω*_max_ ∝ *N*^*−*0.26^ for the stopping frequency.

**Fig. S16:**
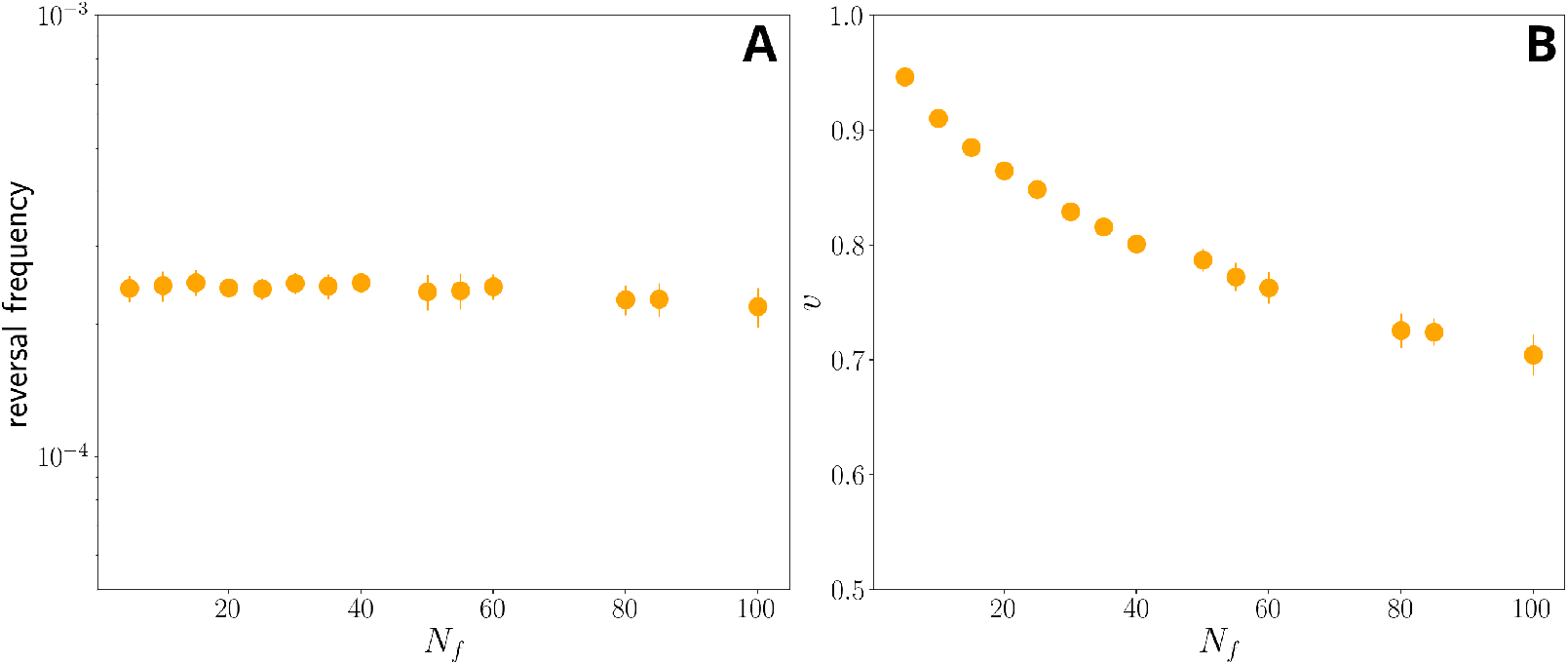
The reversal frequency **(A)** and median filament velocity **(B)** against *N*_*f*_, and *ω*_max_ set using 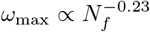 (at fixed *K*_*ω*_=15) for simulations without external field. Notice that, the reversal frequency (left panel) remains roughly constant with *N*_*f*_, while the median velocity decreases. This is a limitation of the model, which does not include any mechanism for reducing filament drag which, as previously mentioned, would rationalize the weak positive correlation between filament velocity and length; however it is consistent with the overall observation that longer filaments tend to be less coordinated.

